# Utilization of an *Artery-on-a-chip* to unravel novel regulators and therapeutic targets in vascular diseases

**DOI:** 10.1101/2022.11.29.517312

**Authors:** Valentina Paloschi, Jessica Pauli, Greg Winski, Zhiyuan Wu, Zhaolong Li, Nadiya Glukha, Nora Hummel, Felix Rogowitz, Sandro Meucci, Lorenzo Botti, Albert Busch, Ekaterina Chernogubova, Hong Jin, Nadja Sachs, Hans-Henning Eckstein, Anne Dueck, Reinier A. Boon, Andreas R. Bausch, Lars Maegdefessel

## Abstract

**Introduction:** Organs-on-chips represent novel *in vitro* models that have the capacity to emulate aspects of human physiology and pathophysiology by incorporating features like tissue-multicellularity and exposure to organ-relevant physical environment. We developed an *artery-on-a-chip* with the objective to recapitulate the structure of the arterial wall composed of intimal and medial layers and the relevant hemodynamic forces that affect luminal cells.

**Results:** By comparing *arteries-on-chips* exposed either to *in vivo*-like shear stress values or kept in static conditions, we identified a panel of novel genes modulated by shear stress. We next measured the expression pattern of shear stress-modulated genes in areas of the vascular tree affected by atherosclerotic plaques and aortic aneurysms, where disease development and progression are induced by alterations of shear stress. We obtained biopsies from patients affected by carotid artery disease (CAD), comprising the atherosclerotic plaque (diseased artery) and the adjacent region (non-diseased artery). From patients with abdominal aortic aneurysms (AAA), we obtained the aneurysmal portion (diseased aorta) and non-dilated adjacent segment (non-diseased aorta). Genes modulated by shear stress followed the same expression pattern in non-diseased segments of human vessels and were expressed by endothelial and smooth muscle cells as evidenced by immunofluorescence analysis and single cell RNA sequencing. Using mice and porcine models of vascular CAD and AAA, we confirmed that shear stress mediated targets are important in discriminating diseased and non-diseased vessel portions *in vivo*. Furthermore, we showed that our *artery-on-a-chip* can serve as a platform for drug-testing. We were able to reproduce the effects of a therapeutic agent previously used in AAA animal models in *artery-on-a-chip* systems and extend our understanding of its therapeutic effect through a multicellular structure.

**Conclusions:** Our novel *in vitro* model is capable of mimicking important physiological aspects of human arteries, such as the response to shear stress, and can further shed light on the mechanism of action of potential therapeutics before they enter the clinical stage.

**Teaser:** The *artery-on-a-chip* is a novel *in vitro* platform that enables the mimicry of human arteries and can be used to gain insights into the development and therapeutic targeting of vascular diseases.

## Introduction

The investigation of etiopathogenetic mechanisms underlying the onset of cardiovascular diseases (CVD), as well as the study of molecular processes involved in disease progression and testing of novel therapies, have all been possible thanks to the availability of a plethora of *in vivo* animal models available. However, the translation of discoveries across species remains challenging when considering the unique nature of the human circulatory system, due to its mechanical, biochemical, and cellular complexities. At the same time, the use of animal models in medical research must adhere to the 3R principles (replacement, reduction, refinement)^1^. In this context, the need to create *in vitro* models capable to mimic key aspects of human physiology/pathophysiology has led to the development of organs-on-chip (OoC), which are new tools to fill the translational gap of “animal-to-human models”^2^. OoC are micro-engineered *in vitro* models of human organs that can recapitulate the minimal functional unit of a tissue/organ and can be used as platforms for pathophysiological studies^3^. They consist of microfluidic cell culture devices containing continuously perfused chambers inhabited by living cells arranged in a dimensional organization that preserves and mimics tissue geometry^4^. In addition, the integration of patient-derived cells and the precise control of the OoC microenvironment enhance our ability to discover novel targets and to perform drug testing using these platforms.

In the present study, we have applied OoC technology to unravel novel factors contributing to two types of cardiovascular diseases: carotid artery disease (CAD) and abdominal aortic aneurysms (AAA). CAD is a common subtype of vascular disease, in which atherosclerosis leads to narrowed arterial lumens and reduced blood flow to the brain, possibly causing transient ischemic attacks (TIA) or strokes^5^. Most commonly, carotid lesions occur at the carotid bifurcation, where a complex and rapidly varying wall shear stress (WSS) distribution is present^6–8^. AAA results from pathological widening of the aortic lumen, an asymptomatic condition that can lead to rupture with poor outcomes (mortality rate >80%)^9,10^. Most of AAAs develop in the infrarenal segment of the aorta, suggesting that specific hemodynamics predispose this portion of the aorta to aneurysmal expansion and rupture^11^. Although CAD and AAA are two distinct vascular diseases^12^, both are caused by pathological remodeling of the vessel wall and influenced by alterations of hemodynamic shear stresses^13^. The endothelial layer, at the interface with blood flow, plays a relevant role in the initial stage of CAD and AAA. Subsequent pathological remodeling of the medial layer populated by smooth muscle cells (SMC) leads to lesion formation and evolution into a diseased vessel wall. Features of CAD include plaque buildup within the arterial wall, disappearance of the intima layer, and increasing inflammation corresponding to disease progression^14,15^. In aneurysms, the vessel wall becomes dysfunctional due to the loss of vascular SMC and destruction of matrix elastic fibers^16,17^.

We herein present the *artery on-a-chip (AoC)* model that mimics the arterial wall structure and captures potential interactions between vascular endothelial cells (EC) constantly subjected to shear stress, and underlying SMC cultured on a layer of extracellular matrix. Moreover, we show how the *AoC* can be used as a molecular target discovery model in CVD research and how these findings translate into the human disease context. Secondly, we provide evidence for the possibility of utilizing the *AoC* as a drug testing model, in which therapeutic agents and their effect in cellular and molecular pathophysiology can be tested.

## Results

### The *AoC*: an *in vitro* model of the arterial wall

The *AoC* consists of a resealable glass chip containing two flow channels separated by an intermediate layer that embeds a porous culture membrane and allows for circulation of two different fluids on each side of the membrane (Fig. **1A**). The intermediate membrane layer creates an interface for co-culture. Primary aortic EC grow on a thin layer of collagen on the flat-side of the membrane, whereas primary aortic SMC are cultured on a fibronectin layer on the well-side of the same membrane (Fig. **1B**). To assess the quality of the coculture, immunofluorescence staining was performed using anti-PECAM and anti-SM22 antibodies, which specifically mark EC and SMC attached at opposing sides of the membrane. An entire scan of the whole membrane was performed (Fig. **1C**). Once cells are confluent, the glass chip is assembled and sealed via a chip holder, which is ultimately connected to a microfluidic pressure controller to perfuse cell-specific culture medium (MFCS-EX from Fluigent, Villejuif, France). The *AoC* is then connected to a MFCS-EX pump and integrated into an incubator. The flow-rate is monitored and mediated externally. Throughout the experiments, the AoC was perfused for 24 hours with a steady flow rate of 1.2 ml/min endowing a constant WSS of 10 dyne/cm^2^ on the EC channel. The flow-rate in the SMC channel is 10 μl/min corresponding to a 0.0021 dyne/cm^2^ WSS, a negligible shear stress that ensures the physiological turnover of nutrients and wastes in accordance with the *in vivo* situation. The flow sensors of the microfluidic platform guarantee a steady flow-rate. The two 2-Switch valves (Fluigent, Villejuif, France) allow the unidirectional recirculation of EC medium between the reservoirs connected to the EC channel (Fig. **1D**).

**Figure 1.**
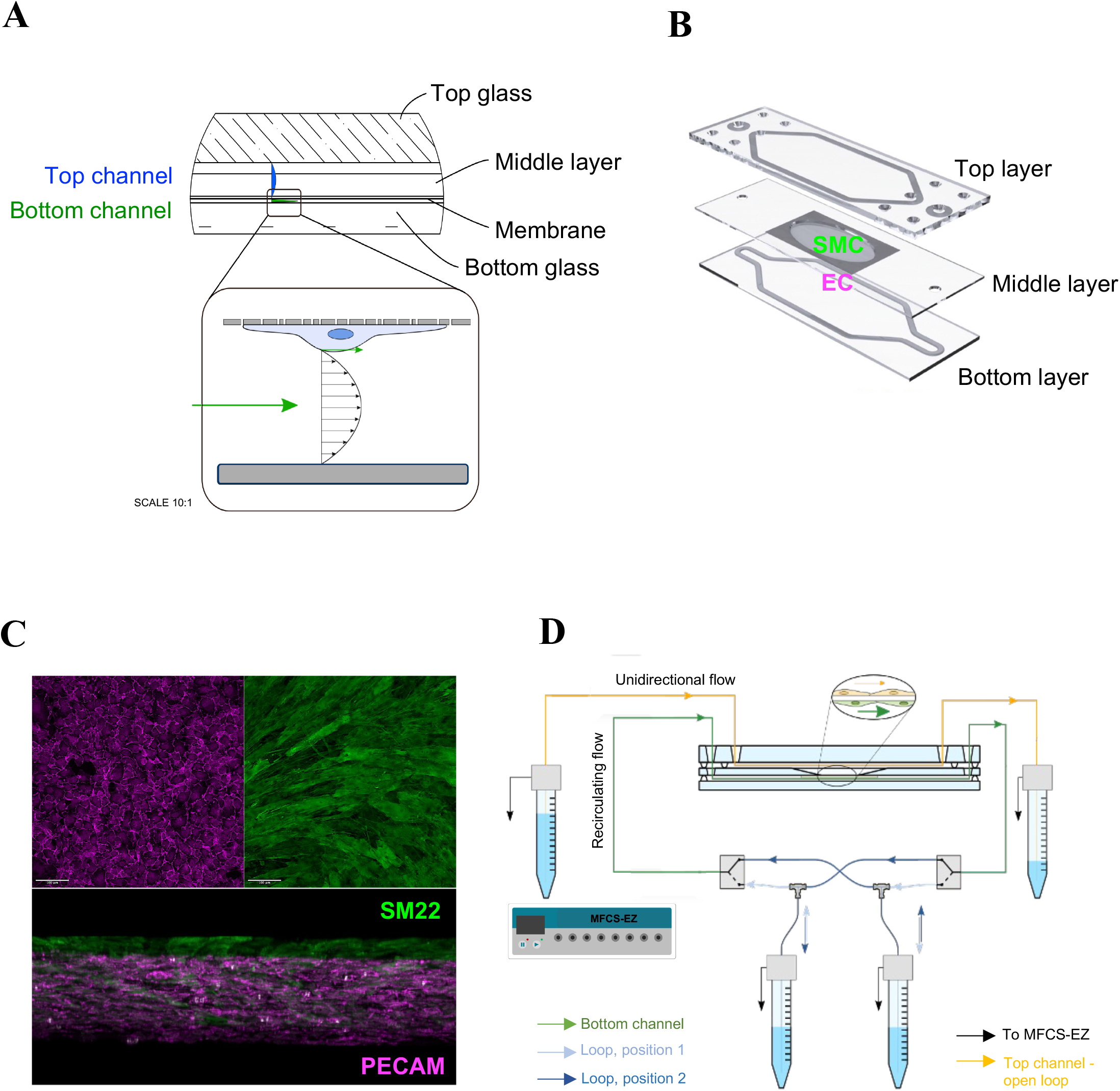
The *AoC* model. (**A**) Cross-sectional design depicting the layers of the *AoC* and the different flow profiles in the top (blue) and bottom (green) channel, respectively. (**B**) The membrane for co-culturing EC and SMC at opposite sides is inserted between the top and bottom layer. (**C**) Immunofluorescence staining of the membrane shows EC and SMC, labelled by the respective cell markers (PECAM for EC and SM22 for SMC). (**D**) Connection of the *AoC* to the microfluidic pump. Arrows indicate the flow direction or the connection to the pressure controller (MFCS-EZ).

Cells were carefully collected at both sites of the membrane, and messenger RNA (mRNA) expression of a panel of marker genes was measured to confirm specific cell identities at the time of cell isolation. Collagen isoforms *COL1A1, COL1A2* and *COL3A1* are expressed by SMC at high levels while barely present in EC. *Vice versa*, von Willebrand factor (*VWF)* and *PECAM* are specifically expressed in EC, and present only at very low levels in SMC (Fig. **2A, B**). Moreover, EC and SMC grown at opposite sides of the membrane had a similar expression pattern profile of cells growing in single culture on regular culture flask (Fig. S1A, B).

**Figure 2.**
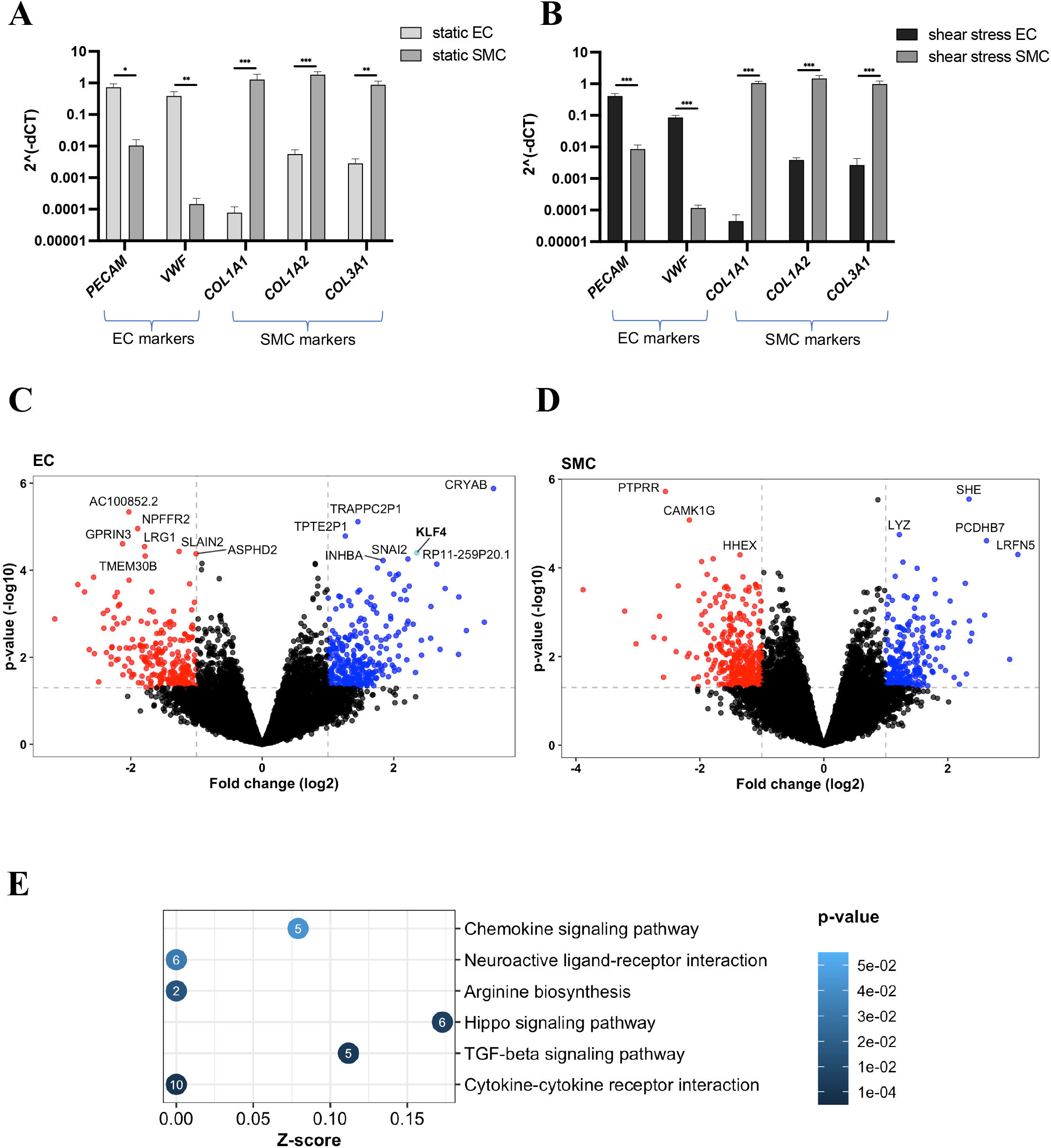
Flow-mediated transcriptomic changes identified in the *AoC*. Gene expression levels measured by quantitative real-time PCR (qRT-PCR) of EC/SMC specific markers performed on cells isolated from shear (**A**) and null stress (static) exposed membranes (**B**). Expression levels are normalized on *RPLPO* levels. Volcano plots depicting EC (**C**) and SMC (**D**), down-(red) and up-(blue) regulated mRNAs in *AoCs* exposed to shear *versus* null stress, as resulted by RNAseq experiments. Differentially expressed genes (DEGs) were identified using a statistical threshold of *P* < 0.01 and fold change ≥ 2. (**E**) Gene-set overrepresentation analysis of KEGG pathway enrichment with enrichment Z-score on the X axis and - log_10_(p-value) on the Y axis. Point size represents pathway size and point color represents Z-score calculated as Z = (S_u_ – S_d_)/√*N*, where S_u_ and S_d_ are the number of significant up- and down-regulated genes in the pathway respectively, and *N* is the total number of genes in the pathway.

### Flow-mediated transcriptomic changes identified in the *AoC*

RNA-seq profiles were generated from 4 *AoC*s exposed to flow while 4 chips were kept in static conditions. Thus, in total 16 samples (4 EC and 4 SMC for each condition) were assessed for transcriptomic changes separately within both cell types in response to flow. Differentially expressed genes (DEGs) were identified using a statistical threshold of *P* < 0.01 and fold change ≥ 2 (Fig. **2C, D**). Kruppel Like Factor 4 (*KLF4*) and 2 (*KLF2*) are transcription factors previously identified as laminar flow inducible elements that play important roles in the regulation of endothelial function^18,19,20^. *KLF4* was found upregulated in the EC of the *AoC* exposed to shear stress, confirming the ability of our system to replicate flow-mediated effects (highlighted in Fig. **2C**). A KEGG pathway overrepresentation analysis based on the significant DEGs identified in EC in response to flow highlighted interesting, enriched pathways (Fig. **2E**). Overrepresentation of DEGs that belong to “cytokine-cytokine receptor interaction” network and the “Hippo signaling pathway”, whose activity strongly relies on mechanical cues in the surrounding microenvironment^21,22^, imply that EC exposed to shear stress are in an activated, stimulus-triggered state. The overrepresentation of DEGs belonging to the arginine biosynthesis pathway, precursor for the synthesis of the vasodilator nitrogen molecule nitric oxide (NO)^23^, was also observed. NO is considered to be an indicator for the integrity of the endothelium^24^. Accordingly, an intact NO release can be regarded as a sign of a healthy endothelium, suggesting that a wall shear stress of 10 dyne/cm^2^ exerts a protective effect on EC.

### *AoC*-generated targets are identified in vascular tissues and associated to the non-diseased status

Next, we evaluated whether the findings observed in the *AoC* had relevance in specific portions of human vessels where disease development and progression are induced by alterations of shear stress. Thus, we assessed human tissue specimens of the Munich Vascular Biobank from patients with CAD as well as AAA^25^. From CAD patients we obtained biopsies comprising the atherosclerotic plaque (diseased artery) and the adjacent region (non-diseased artery, ctrl) (Fig. **3A**). Similarly, from AAA patients we obtained the aneurysmal portion (dilated aorta) and non-dilated adjacent segment (Fig. S2A).

**Figure 3.**
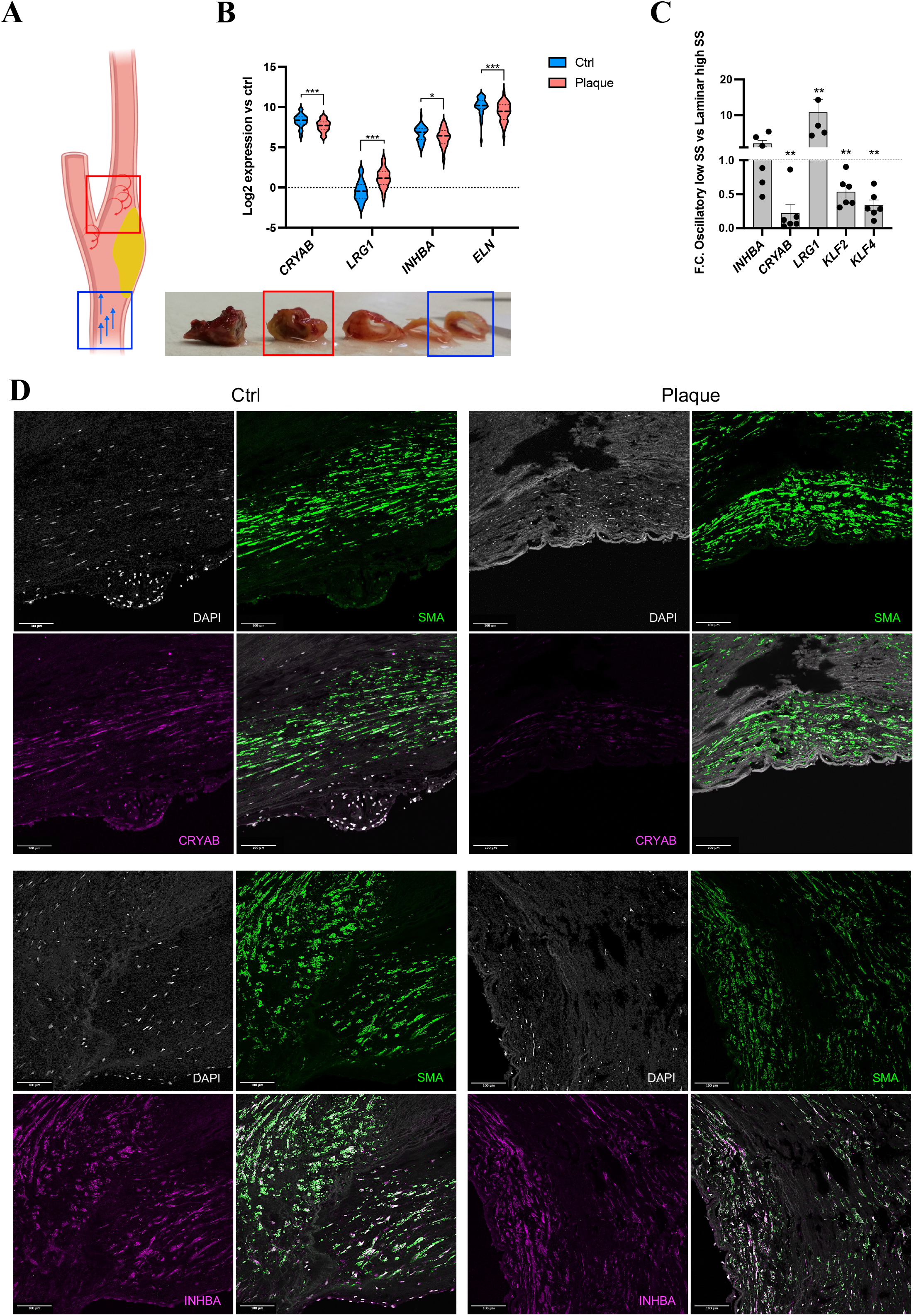
*AoC*-generated targets are identified in human vascular tissues. (**A**) Pictures of human carotid vessels with their respective schematic representations are shown. The biopsies were isolated from the most diseased location (plaque) in the vessels (indicated by red squares) and from the adjacent non-diseased area (internal control indicated by blue squares). Blue arrows show the predicted laminar flow in non-diseased areas, rotating red arrows represents the predicted turbulent flow at the diseased areas. (**B**) Non-diseased (Ctrl) and diseased (Plaque) carotid vessels (n=37) were subjected to RNA-seq. *ELN, CRAYB, INHBA* and *LRG1* expression levels are shown. Statistics: DEGs were determined by paired T-test using a statistical threshold corrected for multiple testing using the false discovery rate (FDR). (*) FDR-adjusted p-value < 0.05; (***) FDR-adjusted p-value < 0.001. (**C**) RNA was extracted from EC exposed for 24h to laminar high stress (12 dyne/cm^2^) and oscillatory low shear stress (2 dyne/cm^2^) and followed by qPCR analysis of novel flow response genes (*CRYAB, INHBA, LRG1, KLF2* and *KLF4*). Paired T-test (n=6) was performed. (**) p < 0.01. (**D**) Double immunofluorescent staining of CRYAB (or INHBA) with SMA and nuclear DAPI in non-disease and diseased human carotid sections. Imaging was carried out with confocal microscopy.

At the carotid plaque site (diseased portion) the shear stress is usually low and oscillatory, while the adjacent, non-diseased portion of the vessel is likely to be subjected to unidirectional flow^6,26^. Similarly, at the aneurysmal site, shear stress alterations are likely present due to dilatation and thrombus formation^11^, while the adjacent, non-dilated aortic segment is subjected to physiological shear stresses. Here, we hypothesized that the two antithetic flow conditions present in diseased and non-diseased vessels, can be matched with the two *AoC* models, namely the flow-exposed, constant shear stress *AoC* and the static, null shear stress *AoC*.

The most significantly upregulated gene in EC in the *AoC* sequencing dataset (flow *vs*. static) is crystallin alpha B (*CRYAB*), a small heat shock protein that primarily binds misfolded proteins to prevent protein aggregation, as well as inhibiting apoptosis and thus contributing to the intracellular architecture^27^. Another flow-induced gene was inhibin subunit beta A (*INHBA*), a member of the TGF-β (transforming growth factor-beta) superfamily of proteins. Both *CRYAB* and *INBHA* appeared significantly upregulated when comparing the non-diseased portion of the carotid vessel to the most advanced, stenotic area of the plaque within the same patient (Fig. **3B**), suggesting that shear stress-induced genes identified in the *AoC* might be of relevance *in vivo*. A reduction of elastin (*ELN*) is a prototypical signature of all diseased vessels^28^, and we sought to monitor its expression pattern in our carotid biopsies (non-disease *vs*. stenosing atherosclerotic plaque of the same patients). As expected, there is a negative correlation between degree of disease and *ELN* expression (Fig. **3B**), highlighting the biological relevance of our cohort. Among the downregulated genes identified from the *AoC* after shear stress exposure (Fig. **2C**), leucine-rich α-2-glycoprotein 1 (*LRG1*) was found significantly upregulated in the diseased portion of carotid artery (Fig. **3B**). Studies have demonstrated that *LRG1* promotes pathogenic neovascularization and angiogenesis through activation of TGFβ1 signaling pathway in EC^29,30^, and that it may also be involved in the inflammation-induced progression of atherosclerosis^31^. The downregulation of *LRG1* strengthens the hypothesis that the applied wall shear stress has a protective effect on EC in our *in vitro* model. *Vice versa, LGR1* upregulation in carotid diseased vessels is concordant with its detrimental role in areas likely affected by low shear stress.

To mimic the hemodynamic conditions in diseased arteries (typically affected by turbulent blood flow and low shear stress), EC were exposed to oscillatory flow with low shear stress (2 dyne/cm^2^) and compared to EC exposed to laminar flow with high shear stress (12 dyne/cm^2^). The low shear oscillatory flow had similar effects as the null shear stress conditions applied to the *AoC* on the expression levels of *CRYAB, LRG1, KLF2* and *KLF4* (Fig. **3C**). This suggests that the flow response identified utilizing the *AoC* setup could be used to discern, within vessels, the diseased from the adjacent non-diseased portions.

In AAA specimens subject to RNA sequencing, paired comparisons of dilated *versus* non-dilated aortic tissues (Fig. S2A) showed a trend of upregulation for both *CRYAB* and *INHBA* in the non-dilated aortic segment (Fig. S2B). *ELN* expression pattern correlated to the aneurysm onset (Fig. S2B). Interestingly, protocadherin beta 7 (*PCDHB7*), a potential calcium-dependent cell-adhesion protein with unknown function in the vasculature system, induced by shear stress in the SMC compartment of *AoCs*, was found to be significantly upregulated in non-dilated aortic specimens (Fig. S2B).

CRYAB and INHBA protein expression was evaluated *via* immunofluorescence in tissue sections obtained from human carotid and aortic biopsies. A loss of protein expression was observed in the diseased carotid vessel as compared to the adjacent non-diseased portion (Fig. **3D**, **4A**). The finding was confirmed in the dilated aneurysmal aorta, as compared to the non-dilated part (Fig. S2C, S3A,B, S4A). CRYAB and INHBA co-localize with typical EC (VWF/PECAM) and SMC (SMA) markers, suggesting that the progression of both, carotid artery disease and AAA, involves EC and SMC cell populations, in which flow alterations augment the expression of responsive genes. Moreover, Western blot analyses preformed on protein lysates from non-diseased/diseased carotid arteries and non-dilated/dilated AAA human specimen reveled a significant downregulation of CRYAB and INHBA in the diseased and dilated vessels, corroborating the immunofluorescence data (Fig. **4B**, Fig. S4B).

**Figure 4.**
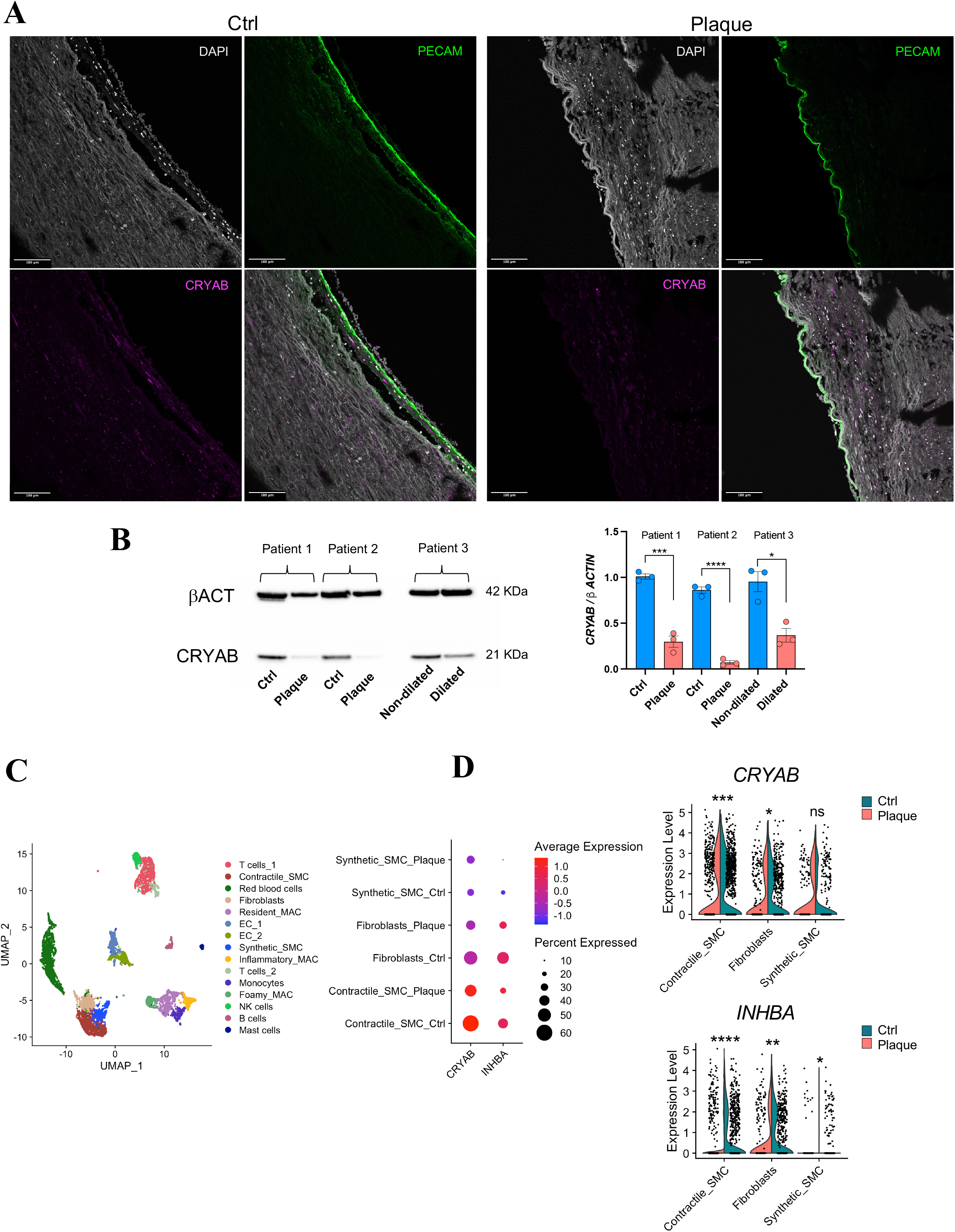
Decreased flow-induced target CRYAB in diseased carotid vessels and aortas. **(A)** Immunofluorescence on non-disease and diseased carotid (plaque) sections showing loss of CRYAB (purple) as well as a decrease degree of co-localization with EC marker, PECAM (green) signal in diseased specimens. Imaging was carried out with confocal microscopy. **(B)** Protein lysates extracted from non-diseased/diseased (plaque) carotid tissues (n=2) and non-dilated/dilated aorta (n=1) were submitted to western blot analysis for the detection of CRYAB. Protein levels are expressed as a ratio to beta actin (βACT). Quantification of the Western Blot was done with Fiji Image J software. Statistics: unpaired T-test (*) p-value < 0.05. **(C)** Uniform Manifold Approximation and Projection (UMAP) plot showing the major cells clusters identified from scRNA-seq performed on human carotid plaques (n=9 patients. Ctrl and Plaque tissues are obtained from each patient). **(D)** Dot plot and violin plot showing the higher expression and significant enrichment of *CRYAB* and *INHBA* in contractile SMC and fibroblast cell clusters originated by Ctrl as compared to Plaque specimens. The FindMarkers function was used to compare the DEGs between the Ctrl and Plaque, by using the default “Wilcoxon Rank Sum test”. (****) p<0.0001, (***) p<0.0002, (**) p<0.0021, (*) p< 0.0332.

### Vascular smooth muscle and endothelial cells are the main cell cluster expressing flow-induced genes in human vascular disease

Single cell RNA sequencing (scRNA-seq) was performed on human carotid plaques (Fig. **4C**, Fig. S5A-C) and human AAA biopsies (Fig. S5D) on samples from the Munich Vascular Biobank. The expression levels of *CRYAB* and *INHBA* were interrogated in both datasets. In the carotid plaque scRNA-seq dataset, *CRYAB* and *INHBA* were found predominantly expressed in the fibroblast-vascular SMC (Fibro-SMC) cluster, (Fig. S5B). Moreover, as the non-diseased vessel (Ctrl) and diseased (Plaque) were independently prepared for individual scRNA-seq datasets from the same patients, *CRYAB* and *INHBA* expression level were compared in Fibro-SMC clusters of the two groups and found more expressed in the cell clusters of non-diseased arteries (Fig. **4D**). Similarly, In the AAA scRNA-seq dataset, *CRYAB* and *INHBA* are enriched in structural cell clusters (Fig. S5E, F) and more highly expressed in the non-dilated SMC cluster compared to the SMC obtained from the dilated aorta (Fig. S5G).

In parallel to scRNA-seq, tissue sections of carotid plaques and control vessels were subjected to hybridization based RNA *in situ* sequencing (HybRISS)^32^, a probe-based targeted amplification detection method, that allows for spatial transcriptomic visualization. From a total number of 98 gene probes, we were able to detect 16.583 transcripts in the control vessel (Fig. S6A), as well as 78.245 and 75.920 in two regions of a carotid plaque (Fig. S6B, C). The latter are located in the plaque shoulder regions, an area where advanced lesions destabilize. Depending on the thickness of the fibrous cap, which confers its stability, a plaque can undergo progressive erosion and rupture, culminating in embolization and potential ischemic forms of stroke^33,34^. In both, control (Fig. **5A**) and plaque sections (Fig. **5B, C**) *CRYAB* and *INHBA* transcripts were detected in cells expressing *MYH11* and *ACTA2* transcripts and, to a lesser extent in von Willebrand factor (*VWF*) positive cells.

**Figure 5.**
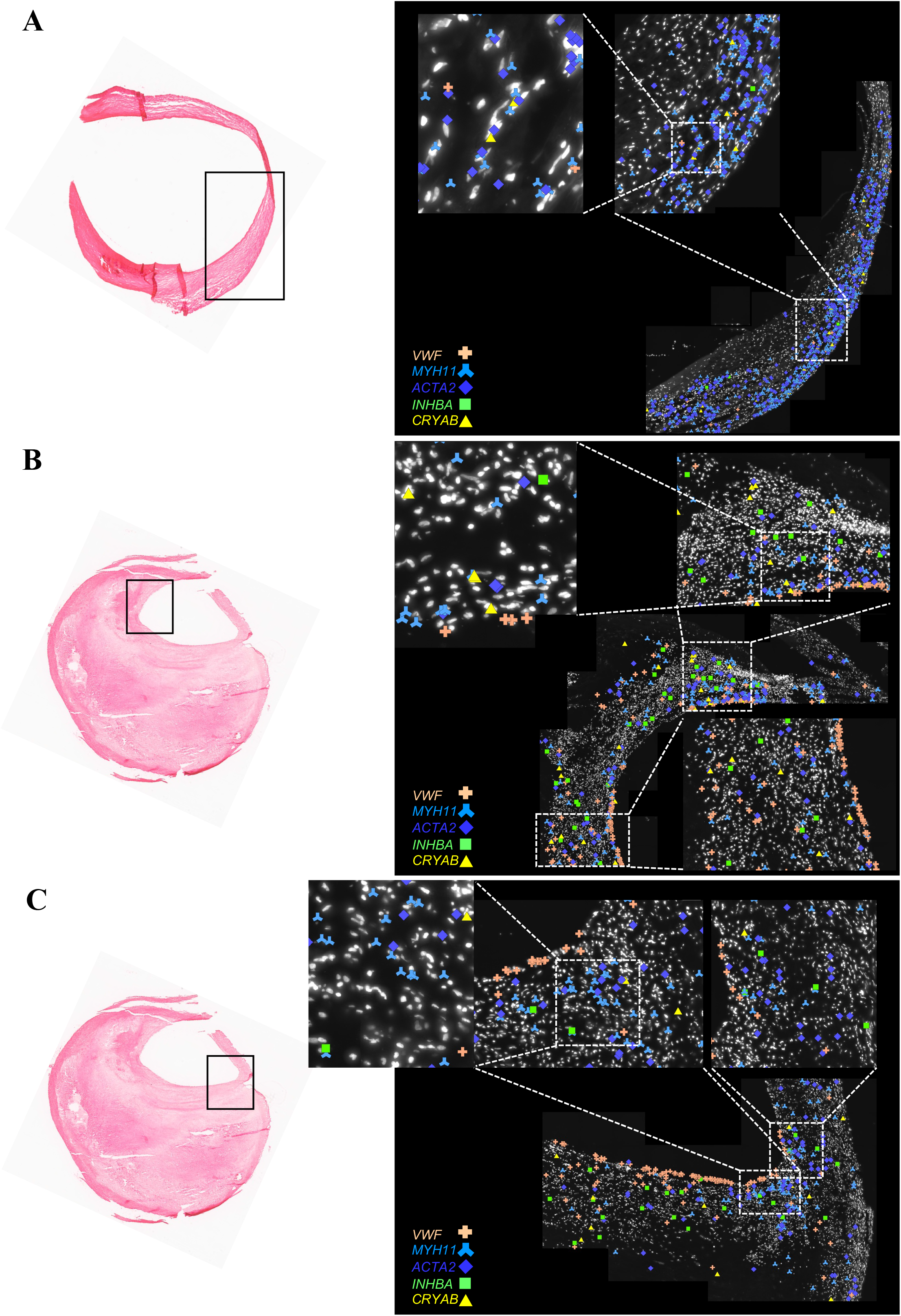
Spatial transcriptomic of flow-induced targets. (**A**) Ctrl (non-diseased carotid vessel) and (**B, C**) carotid Plaque were subjected to HybRISS methodology. *VWF, ACTA2, MYH11, INHBA* and *CRYAB* transcripts were detected in the tissue section and are presented in different colors. The respective processed areas are marked. In the Plaque section two different regions of the fibrous cap were analyzed.

### *AoC*-identified targets in preclinical experimental vascular disease models

Next, we analyzed scRNA-seq data generated from mice subjected to either an inducible plaque rupture model^35^ or to porcine pancreatic elastase (PPE) induced AAA formation^36^ (Fig. **6B, G**). Sham-treated mice were used as appropriate controls. Similar to the human situation of advanced vascular disease we induced an artificial flow disturbance in both preclinical mouse models. This was either provoked *via* ligation/obstruction of the carotid artery (Fig. **6A**), or by subjecting the abdominal aorta to elastase-collagenase enzyme cocktail that destroys the extracellular matrix and therefore also induces disturbed flow in the vessel (Fig. **6F**). Both mouse models can therefore not only be used to investigate the drivers of CAD and AAA disease, but also to investigate flow-mediated transcriptomic changes *in vivo*. scRNA-seq was performed separately on the treated (“diseased”) and sham/non-treated (“ctrl”) mice vessels, which allowed us to compare biologically relevant tissues, similarly to the approach utilized for human tissues.

**Figure 6.**
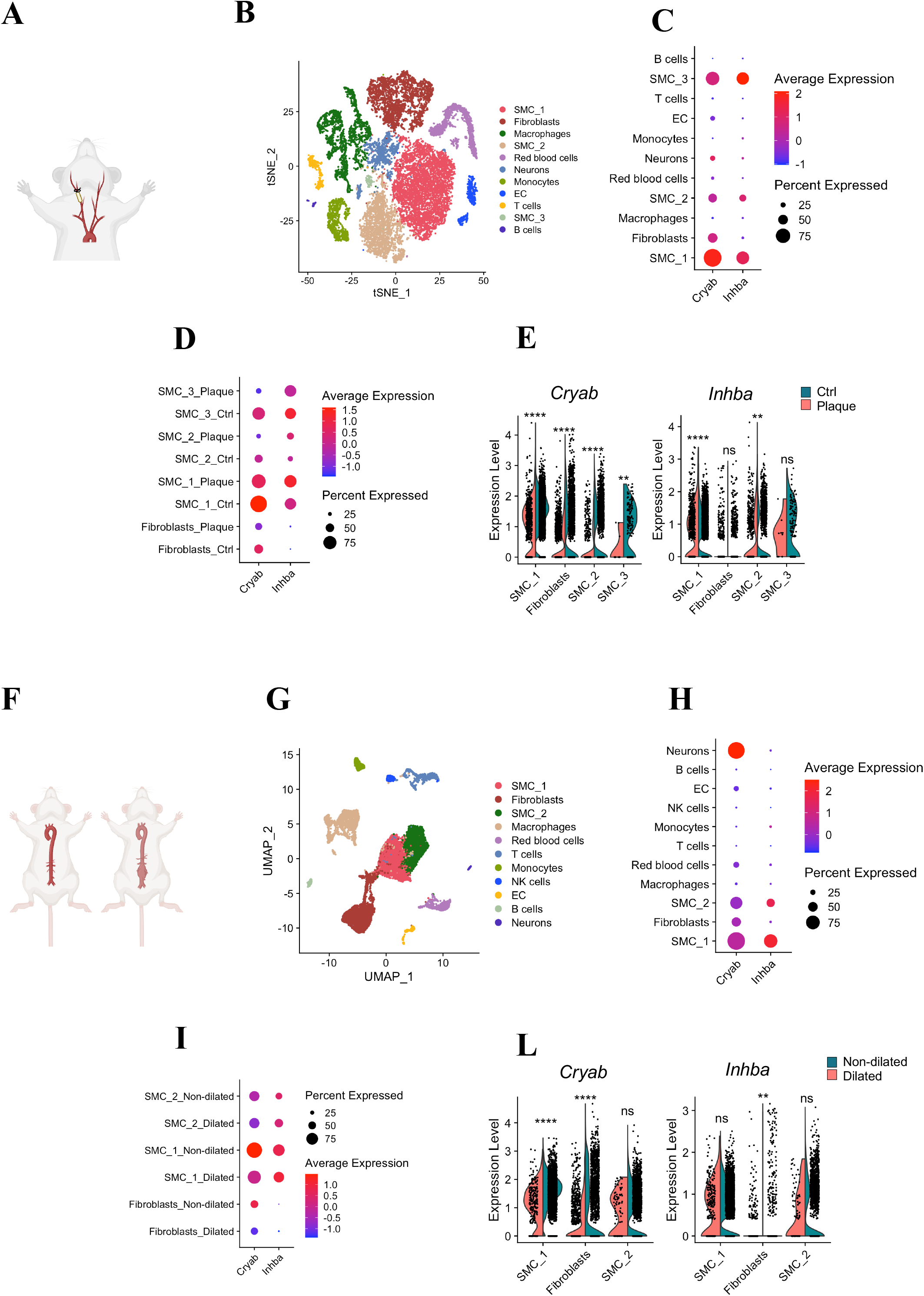
Vascular SMC express flow-induced genes at single-cell level in mice model of CAD and AAA. (**A**) Inducible plaque rupture mouse model: carotid vessels were isolated from control side (left) or subjected to ligation (right). (**B**) t-distributed stochastic neighbor embedding (t-SNE) plot showing the major cells clusters identified from scRNAseq performed on mice subjected to carotid ligation. n=9 in control and carotid plaque group (control and diseased tissues are taken from the same mouse). (**C**) Dot plot showing *Cryab* and *Inhba* enrichment in Fibro-SMC clusters. (**D**) Dot and (**E**) violin plot show the higher expression and significant enrichment of *Cryab* and *Inhba* in control arteries. (**F**) AAA mouse model (PPE infusion) and sham animals (Ctrl). (**G**) UMAP plot showing the major cells clusters identified from scRNAseq performed on aortas isolated from control mice or subjected to the PPE-induced AAA model. In control group n=5, and in AAA group n=6. (**H**) Dot plot showing *Cryab* and *Inhba* mostly expressed in the Fibro-SMC clusters. (**I**) Dot and (**L**) violin plot showing the significant enrichment *Cryab* and *Inhba* in Fibro-SMC clusters of control mice aortas. The FindMarkers function was used to compare the DEGs between the control and carotid plaque/AAA tissue, by using the default “Wilcoxon Rank Sum test”. (****) p<0.0001, (**) p<0.0021.

In both, plaque rupture and AAA mice models, *Cryab* and *Inhba* were found to be mainly expressed by Fibro-SMC clusters (Fig. **6C, H**). Additionally, *Cryab* and *Inhba* expression appear significantly lower in diseased cells as compared to cells from control animals (Fig. **6 D-E, I-L**). On the contrary, *Lrg1*, a gene downregulated in the *AoC* exposed to shear stress and associated with the disease status of vessels, was significantly higher expressed in the EC cluster originating from murine carotid plaques/AAA tissues (Fig. S7A-H).

Furthermore, we obtained non-dilated and dilated portions of a large, preclinical model of AAA disease in Yucatan *LDLR*^*-/-*^ mini-pigs upon local elastase instillation (Fig. S7I) as previously described^37^. Again, we assessed the expression level of flow-induced targets and discovered a significant decrease of *INHBA* in the dilated compared to the non-dilated segment accompanied by the lessening of *ELN* expression in all the tissue pairs (Fig. S7L, M).

### *CRYAB* and *INHBA* inhibition affects SMC homeostasis

In order to gain insights into the functional role of *CRYAB* and *INHBA* in SMC, we silenced their expression in disease-relevant human carotid SMC. The lack of *CRYAB* and *INHBA* in carotid SMC significantly altered the transcriptome profile and resulted in a great number of significantly DEGs (Fig. **7A**, Fig. S8A, D, G). Gene Set Enrichment Analysis (GSEA) was used to associate a disease phenotype to the DEGs after *CRYAB* silencing and highlighted “artery disease” and “vascular disease” as highly significant disease ontologies (Fig. **7B**, Fig. S8B). Moreover, the Human Molecular Signatures Database (MSigDB) hallmark gene sets were employed to summarize well-defined biological states or processes affected by *CRYAB* silencing (Fig. **7C**, Fig. S8C). The epithelial to mesenchymal transition (EMT) process is of particular interest in the context of SMC biology in atherosclerosis as EMT genes were shown to dramatically alter SMC phenotypes contributing to atherosclerotic plaque composition^38^. Similar results were obtained from *INHBA* silencing in carotid SMC with the additional enrichment of immune system disease ontology (Fig. S8D-I). Indeed, *INHBA*, a member of the TGF-β superfamily of cytokines, plays pivotal roles in the regulation of immune responses^39^. Altogether, *CRYAB* and *INHBA*, identified as flow response genes in our *AoC*, seem to have fundamental importance for the homeostasis of SMC (where they are expressed abundantly) during vascular disease development and progression.

**Figure 7.**
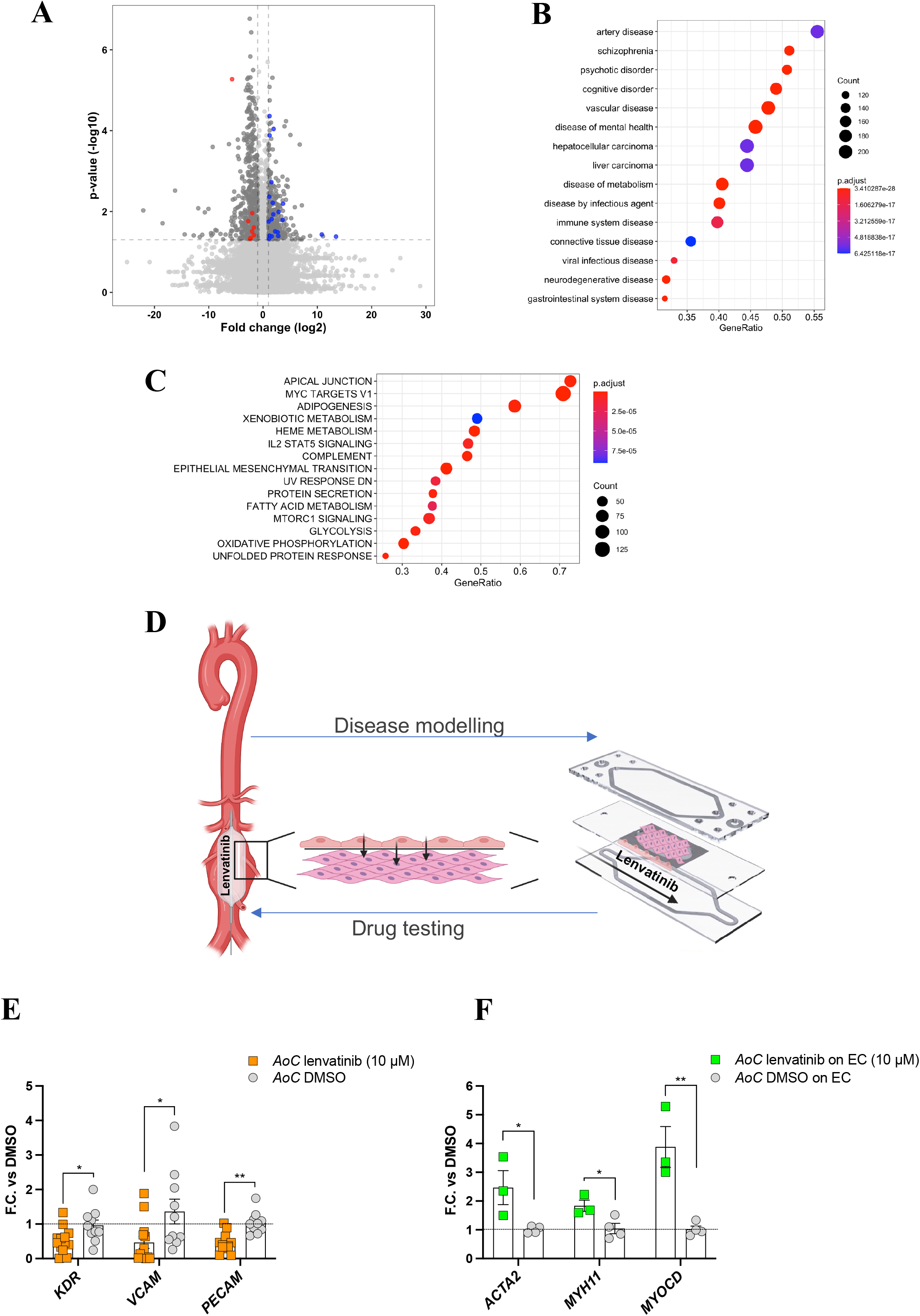
*AoC* as a testing model for therapeutical agent Lenvatinib. (**A**) Volcano-plot with de-regulated genes 48h after transfection with siCRYAB in CaSMC. Highlighted are genes contributing to GSEA score enrichment of Disease Ontology terms “Artery disease” and “Vascular disease”. (**B, C**) Gene Set Enrichment Analysis (GSEA) using Disease Ontology (**B**) and MSigDB Hallmark (**C**) gene sets. (**D**) Scheme representing the utilization of the *AoC* as translational tool. Lenvatinib-coated balloon showed a therapeutical effect in halting aneurysmal progression in a pre-clinical porcine model. AAA disease modelling is partially obtained by *AoC* populated by AAA-derived SMC. Testing lenvatinib on the *AoC* can unravel its mechanism of action. (**E, F**) *AoC*s exposed to lenvatinib or DMSO (ctrl) were submitted to qRT-PCR to monitor gene expression in EC and AAA-derived SMC separately. For EC analysis: DMSO group n=10; Lenvatinib group n=13. For AAA-derived SMC analysis: DMSO group n=4; Lenvatinib group n=3. Statistics: unpaired T-test (*) p-value < 0.05; (**) p-value < 0.01.

### *AoC* as a testing model for therapeutical agent lenvatinib

Next, we wanted to assess whether the *AoC* model can serve as a drug testing device in the context of vascular diseases. This appeared possible and intriguing based on the recirculation process in the endothelial channel, mimicking the intima-blood flow interface conditions observed *in vivo* with possible transcellular diffusion between the cell layers.

We utilized the *AoC* to test the effect of lenvatinib, a tyrosine kinase inhibitor capable of limiting aortic aneurysm expansion *in vivo* by restoring a contractile SMC phenotype in the above-described preclinical murine and porcine models of AAA development and exacerbation^37^ (Fig. **7D**). For these experiments, the EC layer of the *AoC* was exposed to circulating medium containing 10μM of lenvatinib or DMSO (as control), whereas the SMC channel was exposed to SMC medium as described for the original *AoC* set up. The SMC utilized for these experiments were isolated from patients affected by AAA to assess whether it was possible to reproduce the protective effect of lenvatinib determined in previous *in vivo* studies. After 24 hours exposure to lenvatinib or DMSO, cells at the opposite side of the membranes were collected and the gene expression of known lenvatinib targets^37^ were measured. EC responded to lenvatinib with downregulation of *VEGFR2* (*KDR*) and *PECAM* expression, which is consistent with its established inhibitory role in angiogenesis^40^. Moreover, lenvatinib treatment resulted in downregulation of the inflammatory activation marker *VCAM1*^41^ (Fig. **7E**).

Interestingly, patient-derived SMC in co-culture with EC exposed to lenvatinib treatment upregulated genes related to the contractile phenotype (*ACTA2, MHY11, MYOCD*) when compared to SMC co-cultured with EC exposed to DMSO (Fig. **7F**). This protective effect in the context of aneurysm disease recapitulated *in vivo* observations from our recent intervention study, and is an important observation for the execution of future clinical studies in AAA patients.

## Discussion

The *artery-on-a-chip* mimics the architecture of the arterial vessel wall and allows to study changes induced by wall shear stress (WSS) relevant to vascular disease development and progression.

WSS is the frictional force *per* unit area applied by the blood flow on the endothelium and it regulates important physiological blood vessel responses, such as the acute vessel tone regulation and the development of blood vessel structure during embryogenesis, as well as chronic remodeling and progression of vascular wall pathologies^42^. Imaging techniques, like phase contrast MRI, can be used to estimate blood velocity (at several locations within a given cross-sectional vessel area) and compute approximations of velocity gradients near walls^43,44^. Similarly, computational fluid dynamic analyses can be employed to predict WSS values. In a straight vessel, characterized by the Womersley-like flow behavior, vascular EC are continuously exposed to unidirectional high-shear solicitations. As a result, EC are organized in a quiescent, anti-thrombotic monolayer^45^. In contrast, flow conditions at branch points and bifurcations are often characterized by complex secondary vortices and flow instabilities resulting in low and/or oscillatory shear stresses. These solicitations promote EC proliferation and a pro-atherogenic status of the vessel wall^46,47^. Hemodynamic studies have shown that in healthy individuals the mean value of the wall shear stress (time averaged over the cardiac cycle) ranges between 8 and 10 dynes/cm^2^ in the suprarenal aorta^48^ and falls around 8 dynes/cm^2^ in the common carotid artery^49^. In contrast, low values of WSS are predictive of carotid plaque development^26^ and correlate with AAA risk of rupture^11^.

In the present study, we tested our newly assembled *AoC* in physiological shear stress (constant WSS of 10 dynes/cm^2^) *versus* static condition and discovered that it can detect biologically relevant flow-inducible genes. *In vitro* flow-induced expression of *CRYAB* and *INHBA*, was confirmed in non-diseased vessels obtained from patients, both at the RNA and protein level. *INHBA* encodes for the inhibin βA subunit of the homodimer Activin A (βA-βA), a multi-functional cytokine belonging to the TGF-β superfamily that has been shown to inhibit vascular endothelial cell growth and angiogenesis^50^, coherently with a quiescent EC monolayer exposed to physiologically relevant shear stress. The increased expression of *CRYAB* observed both in our *in vitro* model as well as in non-diseased human vessels could also be interpreted as protective, given its function in stabilization of cytoskeleton in response to mechanical stress^51^. These observations may support our hypothesis that the selected WSS of 10 dyne/cm^2^ shows a vaso-protective effect on EC *in vitro*, with the results being translatable to human specimens. CRYAB and INHBA were indeed identified *via* immunofluorescence in the healthy portion of carotid arteries and aortas, and appeared to co-localize with both, EC and SMC markers.

Furthermore, the expression of *CRYAB* and *INHBA* in the vessel wall was confirmed *via* scRNA-seq. Fibro-SMC clusters are the mayor contributor to *CRYAB* and *INHBA* expression in both AAA and carotid data of human, with both transcripts significantly more expressed in Fibro-SMC/SMC clusters of non-diseased compared to diseased vessels. Spatial transcriptomics is a complementary method to scRNA-seq, allowing the visualization of specific transcripts in a tissue section to overcome the bias introduced by digestion protocol typical for scRNA-seq of vascular tissues^52^. This is the first time HybRISS has been performed on vascular tissue and carotid plaque sections. It allowed us to reveal that *CRYAB* and *INHBA* transcripts are found mostly in cells expressing *MYH11* and *ACTA2* transcripts, which again confirms SMC produce these transcripts. ScRNA-seq was also performed in preclinical murine AAA and carotid plaque rupture models, where we had the possibility to obtain also healthy aortas and carotid arteries (sham-operated animals). The comparison between the dataset originating from diseased and healthy murine vasculature showed that *Cryab* and *Inhba* are significantly higher expressed in cell clusters from control aortas and carotids. This evidence contributes to the hypothesis that flow-induced genes are present in control arteries and aortas, whereas a loss of expression is associated with the disease vessel status across different species (humans, mini-pigs and mice). Overall, the loss of *CRYAB* and *INHBA* expression seems to have detrimental effects on the homeostasis of SMC in the arterial wall in human disease and experimental models, while being associated with vascular and immune system dysfunction.

It is intriguing to envision the *AoC* as drug testing tool. Any potential therapeutic agents can be transported directly to the cells *via* circulation, enabling us to determine the effect on the EC layer. On the other hand, the co-cultivation and intercellular communication ensured by the semi-permeable membrane offers the additional possibility to investigate the effect of the drug on the underlying SMC fraction. Administration of lenvatinib to the EC flow channel has most probably generated a signal in the EC, that has been transferred to the underlying patient-derived SMC *via* a paracrine mechanism. By using the *AoC*, we provided evidence for the beneficial effect of lenvatinib in restoring SMC contractility, and gain insights into the potential mechanism of action occurring *in vivo*.

Most of the existing vessel-on-chip models were developed to study endothelium-immune cell interaction, with the aim of gaining insights to the pathogenesis of thrombosis. These chips usually consist of single channel lined with EC and perfused with blood flow but lack the integration of SMC. Models that provide perfused co-culture of EC and SMC are rare^53–55^, but essential for monitoring structural changes occurring within the entire vessel wall. In our *AoC* model, the semi-permeable membrane between EC and SMC creates a spatial separation similar to the basal lamina, which demarcates the tunica intima from the tunica media. Second, the application of physiologically relevant shear stress resembling both the blood flow-intima interactions and the intima-media supply by diffusion, allowed us to more accurately mimic human complexity in our *in vitro* setup.

However, the layered structure of the vessel can only be reproduced to a limited extent by our *in vitro* model. For instance, EC and SMC attached to the semi-permeable membrane are aligned in a parallel fashion given the rectangular structure of the two channels. The membrane in the *AoC* is coated with ECM proteins (fibronectin and collagen), thus ensuring a certain interaction between ECM and cells. However, the three-dimensionality of the structure is limited by the fact that the cells and coating solutions are arranged in contiguous layers. In future developments of the *AoC* model, a circumferential alignment of the SMC and the embedding it in ECM proteins would reflect a more accurate approximation of the physiological architecture of the vessel. Moreover, other physiologically relevant types of flow, such as pulsatile, oscillatory, or vorticose flow, common to the vascular branch points, could be interesting for model enhancement.

The *AoC* represents an original model of the arterial vessel wall by incorporating structural aspects of the vasculature (lumen-intima-media) as well as hemodynamic forces impacting the luminal cells. By increasing the capacity to emulate complex aspects of human vascular biology, the *AoC* could partially close the translational gap between cell culture and animal models to human pathophysiology. With the aid of this system, we were able to characterize novel targets previously unrelated to vascular diseases. We were also able to demonstrate that this model system could be used as a platform for testing novel therapeutic agents and strategies in relevant (patient-derived) cell subtypes. These main characteristics demonstrate the translational potential of the *AoC*. In a time where the 3R criteria must be applied in experimental animal research, organs-on-chips are serving as important research tools in unravelling novel targets and therapeutic potential of drugs to human disease. The system will be of great use for future drug target discovery as well as drug testing studies in CVD research.

Finally, utilization of primary patient-derived cells as well as induced pluripotent stem cells (iPSC) for evaluation of syndromic forms of vascular diseases (*e*.*g*., Marfan Syndrome or Vascular Ehlers-Danlos Syndrome) are promising areas of research for the future development of *AoC*.

## Materials and Methods

### Experimental design: *Artery-on-a-chip*

The device utilized in this study has been designed in conjunction with Micronit Technologies (Enschede, The Netherlands). It consists of a membrane layer and two resealable glass slides that form the top and bottom chambers for the *artery-on-a-chip* (*AoC*) device. The membrane, inserted in a glass or plastic carrier, is made of polyethylene terephthalate (PET) and has a pore size of 0,45 μm and a surface of 1 cm^2^. The bottom side of the membrane has a distance to the bottom glass channel of 0,1 mm and 0,25 mm to the top glass channel. Prior to cell seeding the membrane was coated on the bottom with 0.1 mg/ml collagen rat tail (Corning, USA) and on the top, where it forms a well, with 20 μg/ml of fibronectin (Sigma Aldrich) to ensure cell attachment. Cells utilized on the *AoC* were primary human aortic endothelial cells (EC) and human aortic smooth muscle (SMC), both derived from healthy donors (Cell Application, CA, USA). Patient-derived SMC (described in the following paragraph) were also cultured on the *AoC*. EC and SMC were used between passages 3-7 and cultured in EC and SMC Growth Medium (PeloBiotech, Planegg, Germany), respectively.

First EC are seeded on the bottom side of the membrane at a dilution of 100000 cell/cm^2^ and allowed to attach overnight. On the following day the membrane is flipped, and SMC are seeded (80000 cells/cm^2^) on the top concave side (well). On the third day, the membrane with the two distinct cell layers, is placed between top and bottom glass layers forming two separate flow chambers. The assembled *AoC* is secured in the chip holder, a cell culture platform, with separately controlled fluid-flows above and below. The fluid control is exerted by the connection via perfusion set to a microfluidic pressure-based pump (MFCS-EZ from Fluigent, Villejuif, France). Before the *AoC* connection to flow, reservoirs are filled with cell culture medium and pressurized in order to fill and rinse the tubing. To maintain the desired flow rate at a constant level along the experiment, flow sensors that monitor the flow rate are installed between the reservoirs and the *AoC*. A regulation algorithm is continuously evaluating the current flow rates and adjusting the pressure according to fluctuations in the flow. Flow sensors that monitor the flow rate are installed between the reservoirs and the *AoC*. The fluidic set up constituted by the *AoC*, cell culture medium reservoirs, tubing, and flow sensors are placed in a cell culture incubator (37°C, 5% CO_2_) and allowed to equilibrate to minimize temperature variations. The pump is placed outside the incubator and operated by MAESFLO and MAT (Microfluidic Automation Tool) software. At the end of the flow experiment the *AoC* is disassembled, the membrane is rinsed with PBS (Gibco, Thermo Fisher Scientific) and processed for immunofluorescence staining (see paragraph below) or the harvest of EC and SMC separately. For the latter, the membrane is placed on top of a small petri dish lid with the EC facing the bottom, in contact with EC medium to prevent cells from detaching. The SMC on the concave side of the membrane, facing upwards, are washed with PBS and incubated with 100 μl of 0,25% trypsin-EDTA (Gibco, Thermo Fisher Scientific) for 2 minutes, collected into an Eppendorf tube and centrifuged at 11200 rpm at 4 °C for 6 min followed by pellet resuspension with 200 μl of QIAzol (Qiagen, Hilden, Germany). To isolate EC from the bottom layer, the membrane is flipped, the surface is rinsed with PBS and then scraped with 200 μl of QIAzol to collect as many cells as possible into an Eppendorf tube. Both tubes were stored at -80°C for subsequent RNA isolation.

### Cell culture under flow

EC were seeded in μSlides ^0.2^ / μSlides ^0.6^ (Ibidi, Munich, Germany) and connected to the pump (Ibidi, Munich, Germany) according to the manufacturer’s protocol. EC were exposed for 24 hours to laminar flow providing a shear stress of 12 dyne/cm^2^ or oscillatory flow providing a shear stress of 2 dyne/cm^2^.

### Munich Vascular Biobank

Human aortic aneurysm samples, as well as human carotid artery specimen were collected from the Munich Vascular Biobank, as described previously upon patients’ informed consent^25^. The biobank is approved by the local Hospital Ethics Committee (2799/10, Ethikkommission der Fakultät für Medizin der Technischen Universität München, Munich, Germany) and in accordance with the Declaration of Helsinki. Primary human aortic SMC from patients were isolated from AAA biopsies, harvested during surgical repair and stored in complete DMEM/F12 Medium (Sigma Aldrich) containing 5% Fetal Bovine Serum (Gibco, Thermo Fisher Scientific) and 1% Pen-Strep (Gibco, Thermo Fisher Scientific). The tissue was placed in a sterile petri dish and washed with PBS (Gibco, Thermo Fisher Scientific). Adventitia, neo-Intima and calcifications were removed, and the remaining media was cut into small pieces using a sterile scalpel. The pieces of tissue were placed in digestion medium (1.4mg/ml Collagenase A, Roche, Mannheim, Germany, in complete DMEM/F12 Medium) in a humidified incubator at 37°C and 5% CO2 for 4-6 h. Cells were strained using a 100μm cell strainer to remove debris. After 2 washing steps (centrifuge 400g, 5 min; discard supernatant, re-suspend in 15ml complete DMEM/F12 Medium) cells were re-suspended in 7ml complete DMEM/F12 Medium and placed in a small cell culture flask in a humidified incubator at 37°C and 5% CO2. Upon confluence, cells were stored in liquid nitrogen or processed immediately.

### RNA isolation and RT-qPCR analysis

Human and porcine carotid artery/aortic tissues were cut in ∼50mg pieces on dry ice. Tissue was homogenized in 700μl Qiazol lysis reagent and total RNA was isolated using the miRNeasy Mini Kit (Qiagen, The Netherlands) according to manufacturer’
ss protocol. EC and SMC from *AoC* experiments were collected as described above, and total RNA was isolated using the miRNeasy Micro Kit (Qiagen, Netherlands) according to manufacturer’s protocol. RNA concentration and purity were assessed using the NanoDrop system (Thermo Fisher Scientific). RNA Integrity Number (RIN) was assessed using the RNA Screen Tape (Agilent, USA) and the Agilent TapeStation 4200. Next, first strand cDNA synthesis was performed using the High-Capacity-RNA-to-cDNA Kit (Applied Biosystems, USA), following the manufacturer’s protocol. Quantitative real-time TaqMan PCR was then performed using primers for the genes listed below:

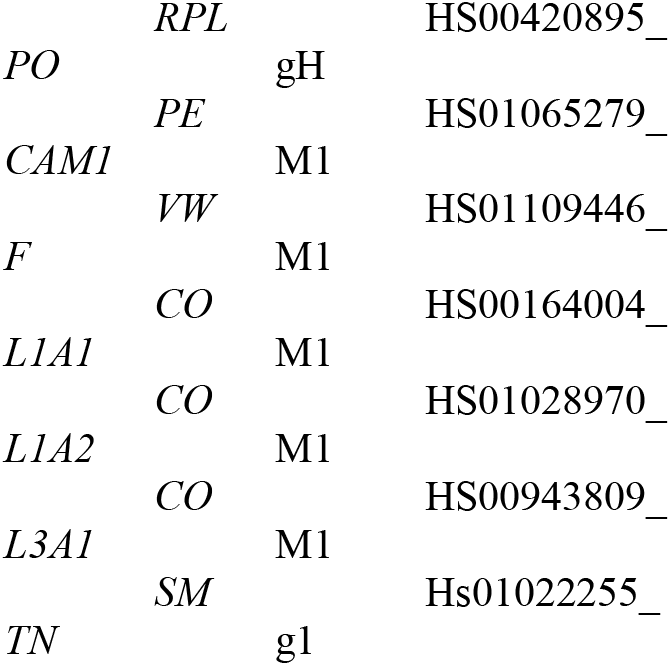
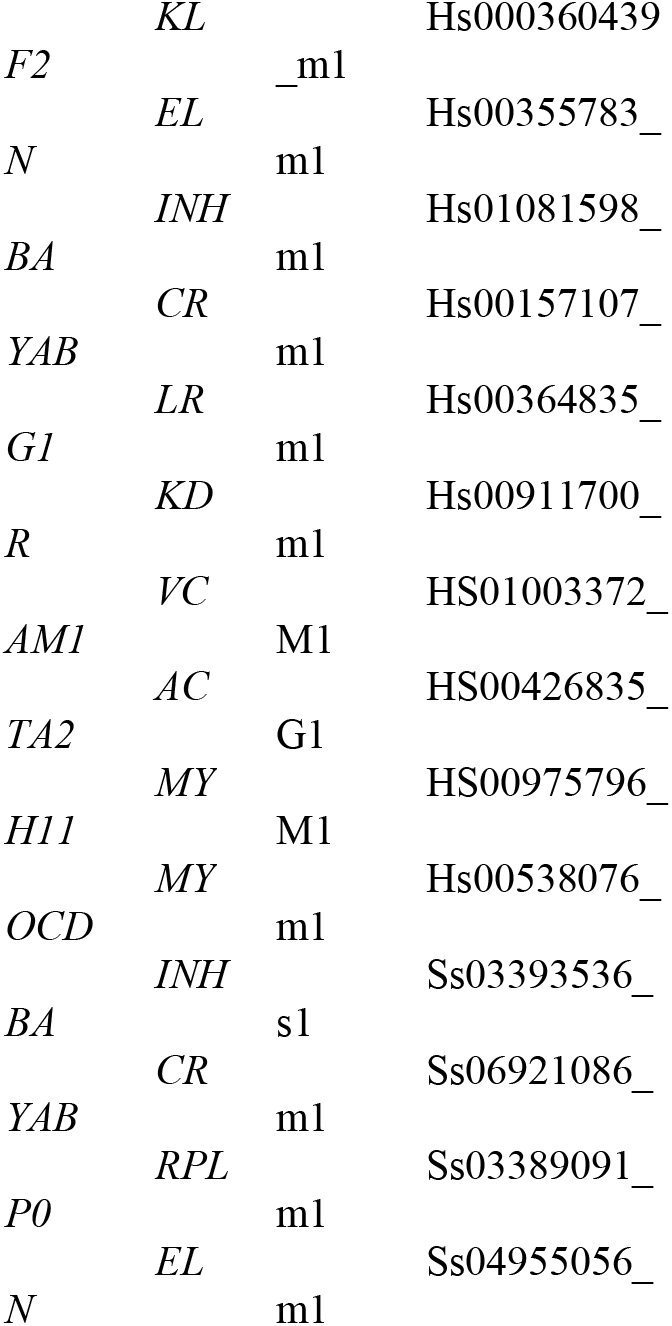

### Cell culture and transfection of human carotid smooth muscle cells

Primary human carotid smooth muscle cells (hCASMC) were obtained from PeloBiotech (#PB-3514-05a) and cultured in Smooth Muscle Cell Growth Medium (PeloBiotech, Munich, Germany), following the manufacturer’s protocol. Transfection of small interference RNA (siRNA) was performed using Lipofectamine RNAiMAX (Thermo Fisher Scientific, Waltham, MA, USA) mixed with anti-*CRYAB*, anti-*INHBA* or scrambled controls (Ambion, California, USA) for a final concentration of 25 nM. Two different timepoints (48hs and 72hs) were investigated. After that, cells were harvested in 700 μl Qiazol and total RNA was isolated using the miRNeasy Micro Kit (Qiagen, Netherlands) according to manufacturer’
ss protocol. RNA concentration and purity were assessed using the NanoDrop system (Thermo Fisher Scientific). RNA Integrity Number (RIN) was assessed using the RNA Screen Tape (Agilent, USA) and the Agilent TapeStation 4200.

### RNA sequencing (RNAseq)

RNA sequencing of EC and SMC exposed to shear/null stress was performed using the IonTorrent Chef System (Thermo Fisher Scientific). A total of 10 ng RNA was reverse-transcribed using Invitrogen Superscript VILO cDNA Synthesis Kit (Thermo Fisher Scientific). Transcriptome libraries were generated using automated library preparation with the Ion AmpliSeq Transcriptome Human Gene Expression Kit (Thermo Fisher Scientific). The transcriptome libraries were barcoded, templated and sequenced on the Ion Chef and Ion GeneStudio S5 Prime systems using Ion 550 Kit-Chef and Ion 550 Chip Kit, as one 16-plex library pool. The quality of the raw expression data was evaluated by Fios Genomics (Edinburgh, UK) using several automated outlier tests. One sample (EC exposed to flow) failed two of these outlier tests and was therefore excluded from subsequent analysis. Expression data was provided as a count matrix for 55,765 features in each sample. 33,357 features with a maximum read count less than or equal to zero across all samples were excluded from analysis. Normalization was carried out using trimmed mean of M-values normalization and expression values were transformed using voom.

Extracted RNA from non-diseased (ctrl) and diseased (plaque) tissue biopsies obtained from 37 patients undergoing carotid endarterectomy as well as from non-dilated and dilated tissue biopsies from 7 AAA patients, was used to generate libraries that where sequenced *via* the Illumina NovaSeq platform. (NovaSeq6000, Illumina, San Diego, USA). RNA was purified from total RNA using poly-T oligo-attached magnetic beads. After fragmentation, the first strand cDNA was synthesized using TruSeq stranded total RNA kit and TruSeq stranded mRNA kit (Illumina, San Diego, USA), followed by the second strand cDNA synthesis using dUTP. The directional library was ready after end repair, A-tailing, adapter ligation, size selection, amplification, and purification. The library was checked with Qubit and real-time PCR for quantification and Bioanalyzer for size distribution detection. Raw data (raw reads) of FASTQ format were analyzed by Fios Genomics (Edinburgh, UK). Quality control (QC) was performed on the raw FASTQ files, and the alignment statistics generated by the STAR aligner during the data processing protocol. All samples exhibited high quality and consistent sequence length distributions, so they were all included in subsequent analyses (Fios Genomics, Edinburgh, UK). RNA extracted from human carotid SMC silenced for *CRAYB* and *INHBA* (and respective controls) was subjected to sequencing the Illumina NovaSeq6000 platform at Novogene (Cambridge, UK).

### Tissue immunofluorescence staining

#### Carotid arteries

3μm sections of formalin fixed and paraffin-embedded (FFPE) human carotid artery samples from the Munich Vascular Biobank^25^ were mounted on 0.1% poly-L-lysine (Sigma-Aldrich, St. Louis, MO, USA) pre-coated SuperFrost Plus slides (Thermo Fisher Scientific, Waltham, MA, USA) and dried over night at 56°C. Sections were dewaxed and heat mediated antigen retrieval was performed using a pressure cooker with 10nM citrate buffer (distilled water with citric acid monohydrate, pH 6.0). Endogenous peroxidase activity was blocked with 3% hydrogen peroxide. Two antibodies were applied in sequence. For each antibody, samples were blocked for 1h (5% horse serum, 1%BSA, 0,5% Triton-X100) and incubated with primary antibodies diluted in 5% horse serum overnight. The appropriate secondary antibody was added for 1h (in 5% horse serum) on the next day. Autofluorescence quenching and counterstaining with DAPI were performed. Images were acquired with an Olympus FLUOVIEW FV3000 (Olympus, Tokyo, Japan) confocal microscope.

#### Aortic aneurysms

Aortic aneurysm tissue was harvested during open repair in our clinical Department for Vascular and Endovascular Surgery (Klinikum rechts der Isar) in Munich. Tissue was placed in RNAlater (Sigma-Aldrich, St. Louis, MO, USA) on ice until further processing. The tissue was cut in several pieces and the most morphologically relevant pieces were snap frozen in liquid nitrogen and immediately embedded in Tissue-Tek O.C.T. (Optimal Cutting Temperature) Compound. OCT embedded tissues were stored at -80°C until further use. Ten μm sections of OCT embedded samples were mounted on SuperFrost Plus slides (Thermo Fisher Scientific, Waltham, MA, USA) and stored at -80°C until further use. Sections were fixed in ice cold acetone for 5 min. Endogenous peroxidase activity was blocked with 3% hydrogen peroxide. Blocking, subsequent antibody staining, autofluorescence quenching, counterstaining with DAPI and imaging was performed as described above for carotid artery FFPE tissues.

##### Antibodies

CRYAB (ab13496, abcam) 1:50

INHBA (ab56057, abcam) 1:25

VWF (ab6994, abcam) 1:3000

PECAM (ab9498) 1:50

SMA (ab7817, abcam) 1:200

SMA (ab5694, abcam) 1:200

Alexa488 anti rabbit (A11034) 1:400

Alexa488 anti mouse (A11001) 1:400

Alexa647 anti rabbit (A21245) 1:400

Alexa647 anti mouse (A21236) 1:400

##### Western blot analysis

Fresh frozen human aortic aneurysm and carotid artery tissue was cut in ∼50mg pieces on dry ice. Tissue was homogenized in 200μl Tissue Extraction Reagent I (Thermo Fisher, USA). After homogenization with the Bio-Gen PRO200 Homogenizer and Multi-Gen 7XL Probes (Pro Scientific, USA), samples were centrifuged for 20 min, 14.000 rpm at 4°C and the supernatant was frozen down at - 80°C. Total protein concentration was measured using the Pierce BCA Protein Assay Kit (Thermo Fisher) following manufacturer’
ss protocol. 10μg of protein from each sample was denatured and reduced at 70°C for 10min, then separated in a Bolt 4-12% Bis-Tris Plus Gel (Thermo Fisher) and transferred onto Trans-Blot Turbo Mini-Size LF-PVDF Membranes (BioRad, Hercules, CA, USA). The blots were blocked with 5% milk in Tris-buffered saline+0.1% Tween-20 for 1h, followed by overnight incubation with the primary antibody against INHBA/CRYAB/βACT in TBS-T+5%milk. After washing with TBS-T, blots were developed with horseradish peroxidase (HRP)-conjugated secondary antibody (Abcam, Cambridge, UK) for 1 h in combination with ECL (GE Healthcare, Chicago, IL, USA). Image detection was performed with C600 Azure Biosystems Imager (Biozym)/ChemiDoc XRS System (BioRad, Hercules, CA, USA). Image quantification was done using ImageJ software.

##### Antibodies

CRYAB (ab13496, abcam) 1:1000

INHBA (ab56057, abcam) 1:1000

β-ACTIN (A5316, Sigma Aldrich) 1:8000

HRP-conjugated secondary (ab205718, ab205719, abcam) 1:10000

### Single cell RNA sequencing (scRNA-seq)

#### Human vessel dissociation

Human carotid or abdominal arteries were harvested during carotid endarterectomy or open repair in our Department of Vascular and Endovascular Surgery (Klinikum rechts der Isar, TUM). The biopsies were minced and digested using the Multi Tissue Dissociation Kit 2 (Miltenyi Biotech, 130-110-203), GentleMACS Dissociator (Miltenyi Biotech, 130-093-235), GentleMACS C tubes (Miltenyi Biotech, 130-096-334) and the 37C_Mulit_G program, all according to the manufacturer’
ss instructions. The cell suspension was strained (70 μm, 40 μm) and Dead Cell Removal (Miltenyi Biotech, 130-090-101) using MS Columns (Miltenyi Biotech, 130-042-201) was performed was performed. Cells were resuspended in PBS + 0,04% BSA.

#### Mouse vessel dissociation

Mouse carotid or abdominal arteries were harvested in Stockholm. The treated (Carotid Plaque Rupture Model n=9 or PPE-AAA n=6) and untreated (carotid artery n=9 or aorta n=5) tissues were transported in PBS on ice to the laboratory. The tissues were cut into small pieces with a surgical scissor and digested using the following enzymes: Liberase 4U/ml (Sigma Aldrich, 5401127001), Hyaluronidase 60 U/ml (Sigma Aldrich, H3506-100MG) and DNAse 120 U/ml (Thermo Fisher, 18047019) in RPMI Medium + 10%FBS at 37°C with agitation for 45 min. The cell suspension was strained (70 μm, 40 μm) and Dead Cell Removal (Miltenyi Biotech, 130-090-101) using MS Columns (Miltenyi Biotech, 130-042-201) was performed. Cells were resuspended in PBS + 0,04% BSA.

#### Single-cell capture and library preparation

Cells were loaded into a 10x Genomics microfluidics Chip G and encapsulated with barcoded oligo-dT-containing gel beads using the 10x Genomics Chromium Controller. Gel Beads-in-emulsion (GEM) cleanup, cDNA Amplification and 3’
sGene Expression Library Construction was performed according to the manufacturer’s instructions (CG000204 Rev D).

#### RNA sequencing

Human: Libraries from individual samples were multiplexed into one lane before sequencing on an Illumina NovaSeq6000 instrument. Mouse: Libraries from individual samples were loaded on an Illumina NovaSeq with 2×150 paired-end kits at Novogene.

#### ScRNA-seq analysis

ScRNA-seq analyses were performed according to Seurat instructions (version 4.0.2) in R (version 4.0.3). Genes were excluded for downstream analysis if expressed in fewer than 5 cells. Cells would also be filtered out in each Seurat object if they fulfilled the following conditions: maximum mitochondrial reads <15%, maximum UMIs fewer than 20,000, maximum genes between 100 and 2000-4000. CellCycleScoring function in Seurat was applied to obtain the scores of cell cycle phases including S phase and G2M phase. SCTransform normalization workflow was adopted to mitigate possible technically driven or other variations, in which mitochondrial genes and cell cycle phase were regressed. T-SNE (t-Distributed Stochastic Neighbor Embedding) was used to convert cells into a two-dimensional map. FindAllMarkers function was performed to detect the main features of each cluster with default parameters. Differentially expressed genes (DEGs) between two groups were obtained by implementing the FindMarkers function in Seurat with default methods and parameters.

#### HybRISS (Hybridization-based RNA in situ Sequencing)

Carotid artery tissue specimen was harvested during carotid endarterectomy in our clinical department for Vascular and Endovascular Surgery (Klinikum rechts der Isar, TUM) in Munich. Tissue was placed in RNAlater (Sigma-Aldrich, St. Louis, MO, USA) on ice until further processing. The tissue was cut in several pieces and the morphological most relevant pieces were snap frozen in liquid nitrogen and immediately embedded in Tissue-Tek O.C.T. (Optimal Cutting Temperature) Compound. OCT embedded tissues were stored at minus 80°C until further use. Ten μm sections were cut with a cryotome and placed on SuperFrost Plus slides (Thermo Fisher Scientific, Waltham, MA, USA). Mounted slides were stored at -80°C until the experiment was performed. Probe-design and the in-situ Sequencing was performed as described previously^32,56^.

### Statistical analysis

Differences in RNA expression, measured by qPCR, were calculated as fold change versus control using the mean ΔCt (defined as Ct^target RNA^-Ct^endogenous control^) within groups and compared using Student’s *t* test. Two-tailed p-values were calculated. For differentially expressed gene analysis of the *AoC* experiments (shear stress vs. null shear), a statistical threshold of p-value < 0.01 with fold change ≥ 2. Significant up- and down-regulated genes (at p-value < 0.01 and fold change ≥ 2) in the “Endothelial Cells (Flow vs. Static)” comparison were mapped to 182 genes and assessed for KEGG pathway enrichment. For RNAseq of human carotid tissues a paired comparison between non-diseased (ctrl) and diseased (plaque) for each patient was performed (37 pairs in total). DEGs were determined using a statistical threshold corrected for multiple testing using the false discovery rate (FDR) adjustment (FDR-adjusted P < 0.05; fold change ≥ 2). For the analysis of transcriptomic changes between non-dilated and dilated samples from patients with AAA, a paired comparison was performed and a significance threshold of unadjusted p-value < 0.01 was applied. For silenced human carotid SMC, differential expression analysis of two conditions/groups was performed using the Ballgown R package and DEG were calculated using FDR adjustment (FDR-adjusted P < 0.05) (Novogene, Cambridge, UK). Differential expression of genes was further analyzed through Gene Set Enrichment Analysis^57^ using clusterProfiler R package together with Disease Ontology^58^ and Molecular Signatures Database Hallmarks^59^ gene sets.

## Acknowledgments

We acknowledge the support of Dr. Chika Okota and Prof. Mats Nilsson from the *in-situ* sequencing facility at SciLife Lab (Stockholm, Sweden) for performing spatial transcriptomics on our human cohort of carotid plaques.

## Author contributions

Conceptualization: VP, LM

Methodology: VP, JP, NG, NH, FR, SM, EC, HJ

Investigation: VP, JP, GW, ZW, ZL

Visualization: NS, HHE, AB, ARB

Supervision: LM

Writing—original draft: VP, LM

Writing—review & editing: VP, LM, LB, RAB

## Data and materials availability

All data and materials used in the analyses are available.

