## Supplementary Figures for "Utilization of an *Artery-on-a-chip* to unravel novel regulators and therapeutic targets in vascular diseases"

### Fig. S1

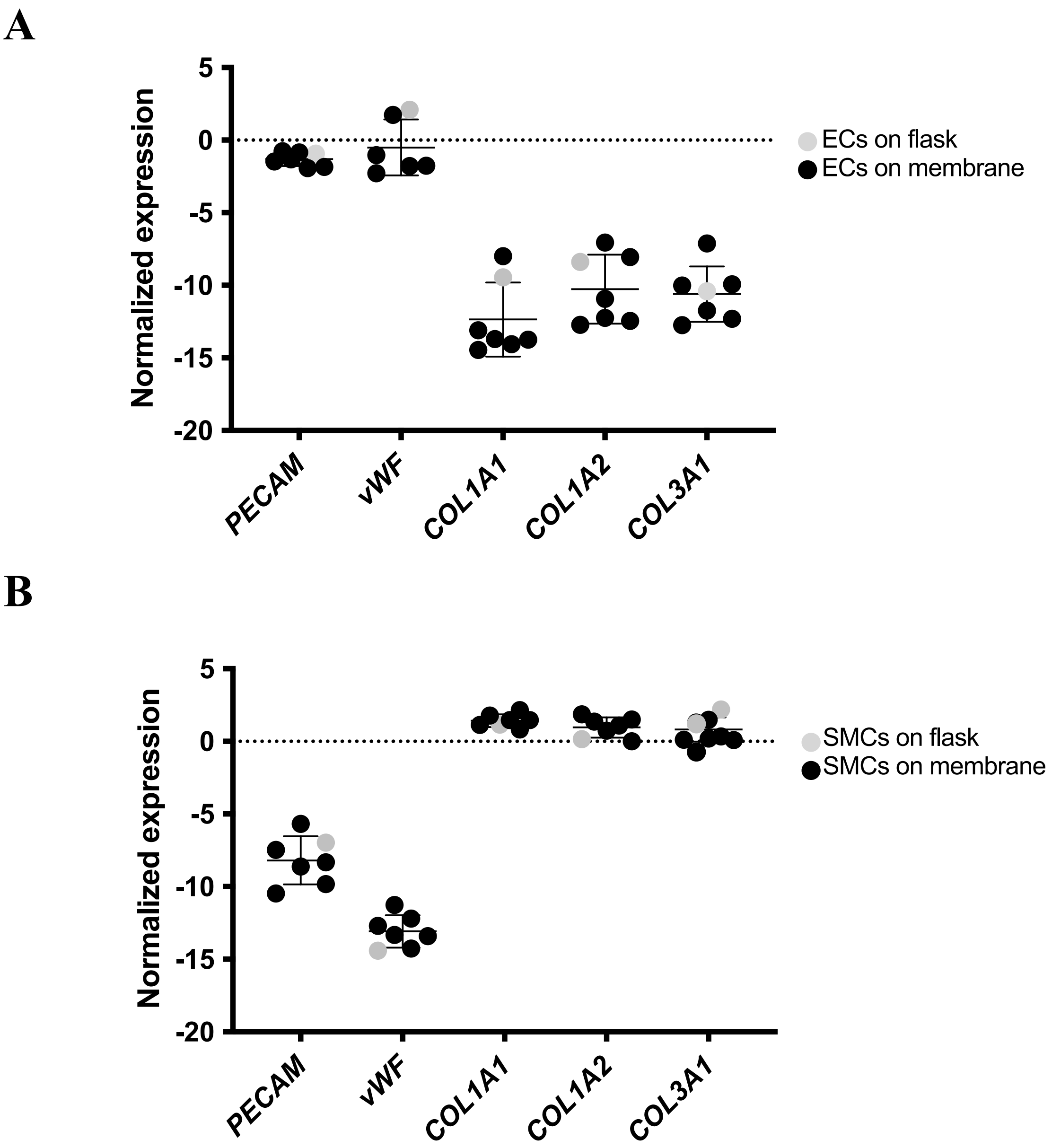


**Figure S1. Comparison of gene expression markers in EC and SMC cultivated on flask or the *AoC* membrane layer.** Distribution of delta CT cycles (normalized on RPLPO) of typical EC (A) and SMC (B) gene markers analyzed from EC and SMC either cultivated on regular cell culture flasks or PET membranes.

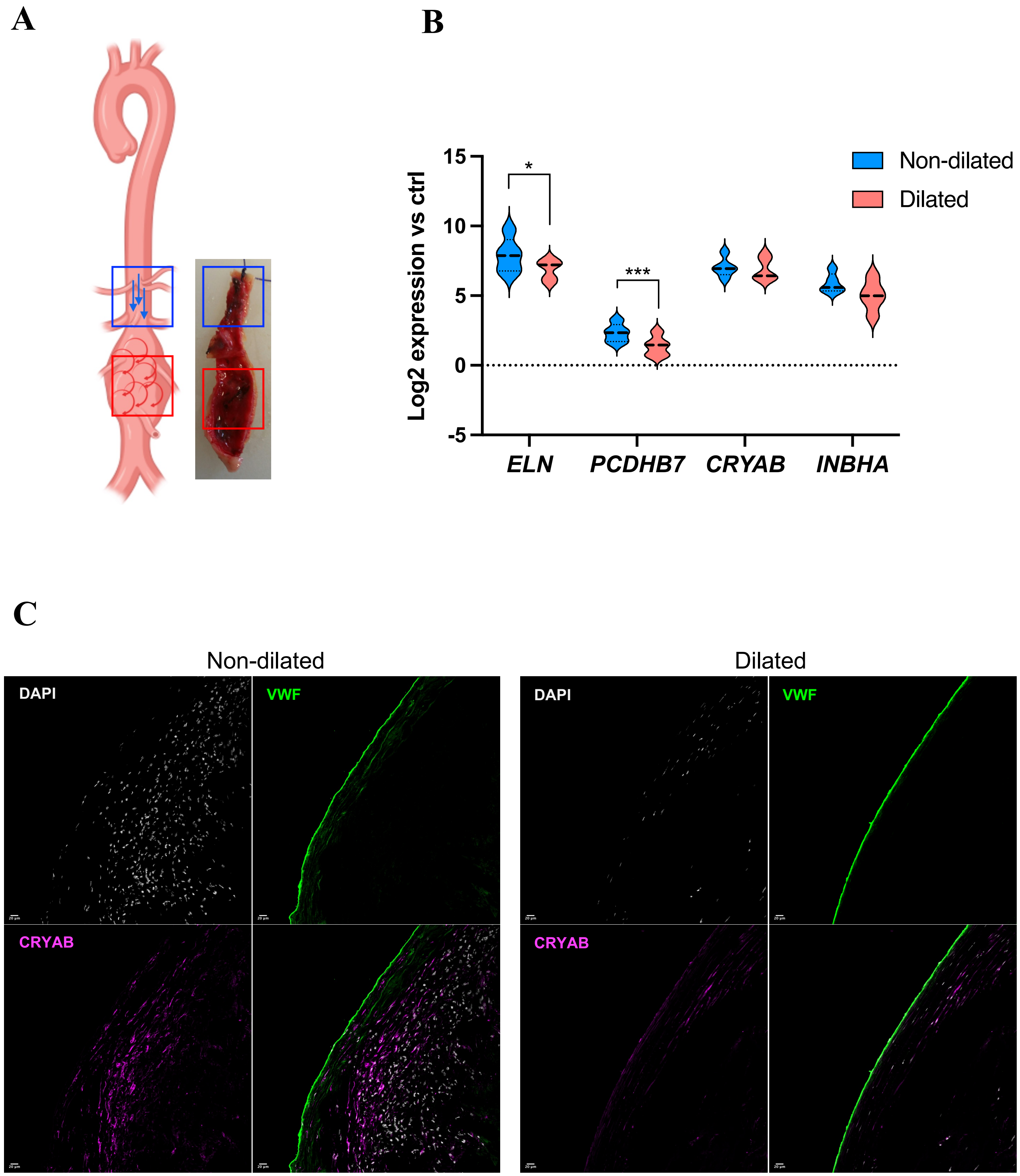


**Figure S2. *AoC*-generated targets are present in human aortas of AAA patients.** (**A**) Pictures of human AAA with their respective schematic representations are shown. The biopsies were isolated from the aneurysmatic area (indicated by red squares) and from the adjacent non-dilated area (internal control, indicated by blue squares). Blue arrows show the predicted laminar flow in non-dilated areas, rotating red arrows represents the predicted turbulent flow at the aneurysmatic areas. AAA specimens from non-dilated (blue square) and dilated (red square) segments were subjected to RNAseq (n=7). (**B**) Log2 expression of *ELN, PCDHB7, CRYAB* and *INHBA*. The latter show non-significant differences between non-dilated/dilated tissue portions. Statistics: paired T-test (*) p-value < 0.05; (***) p-value < 0.001. (**C**) Immunofluorescence on non-dilated and dilated aortic sections showing loss of CRYAB (purple) as well as decrease degree co- localization with VWF (green).

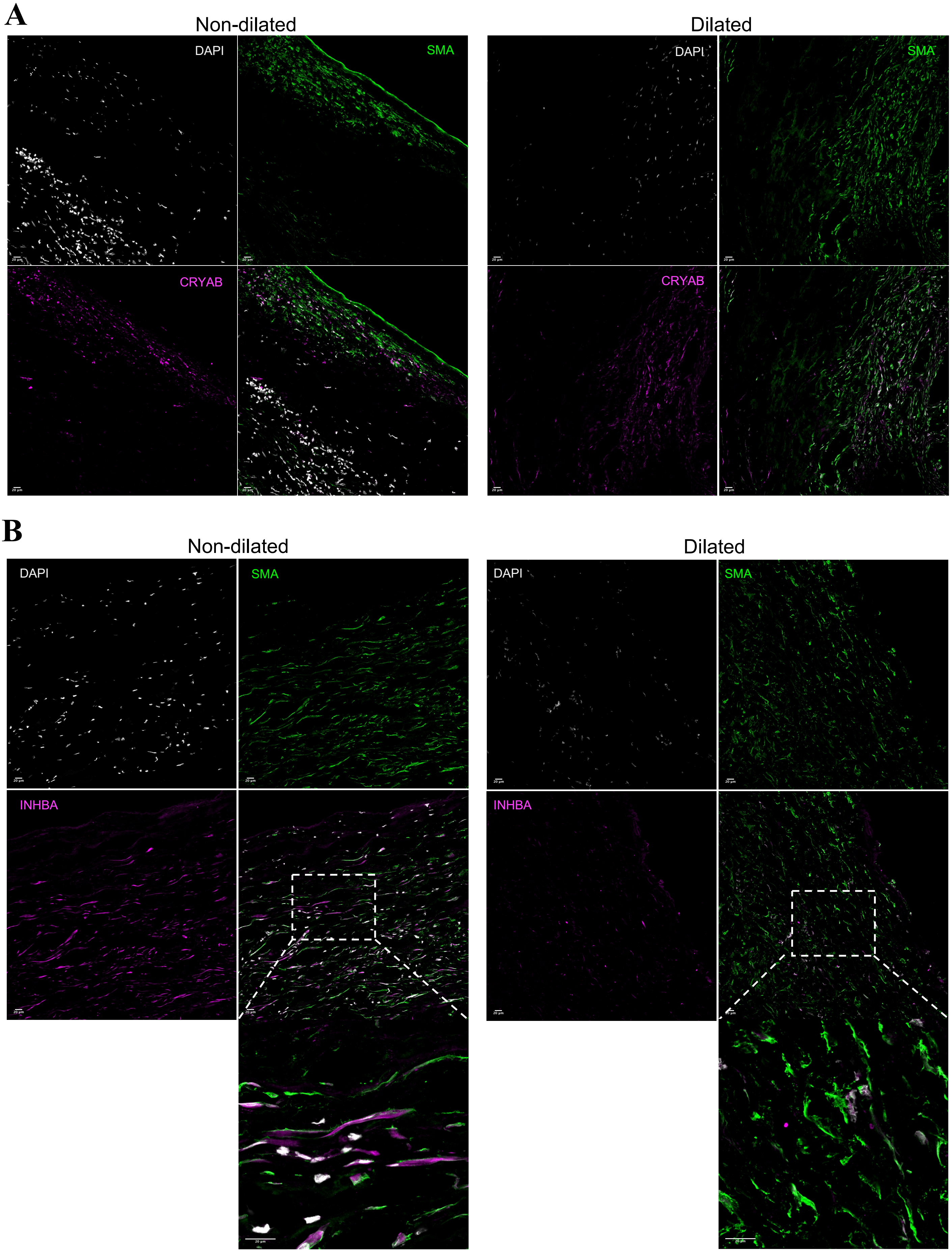


**Figure S3. Decrease amount of CRYAB and INHBA in AAA.** (**A**) Immunofluorescence on non-dilated and dilated aortic sections showing loss of CRYAB (purple) as well as decrease degree co- localization with SMA (green). (**B**) Loss of INHBA (purple) as well as decrease degree of co- localization with SMA (green) in aneurysmal specimens. Zoomed-in images are taken in correspondence to the highlighted white squares. Imaging was carried out with confocal microscopy.

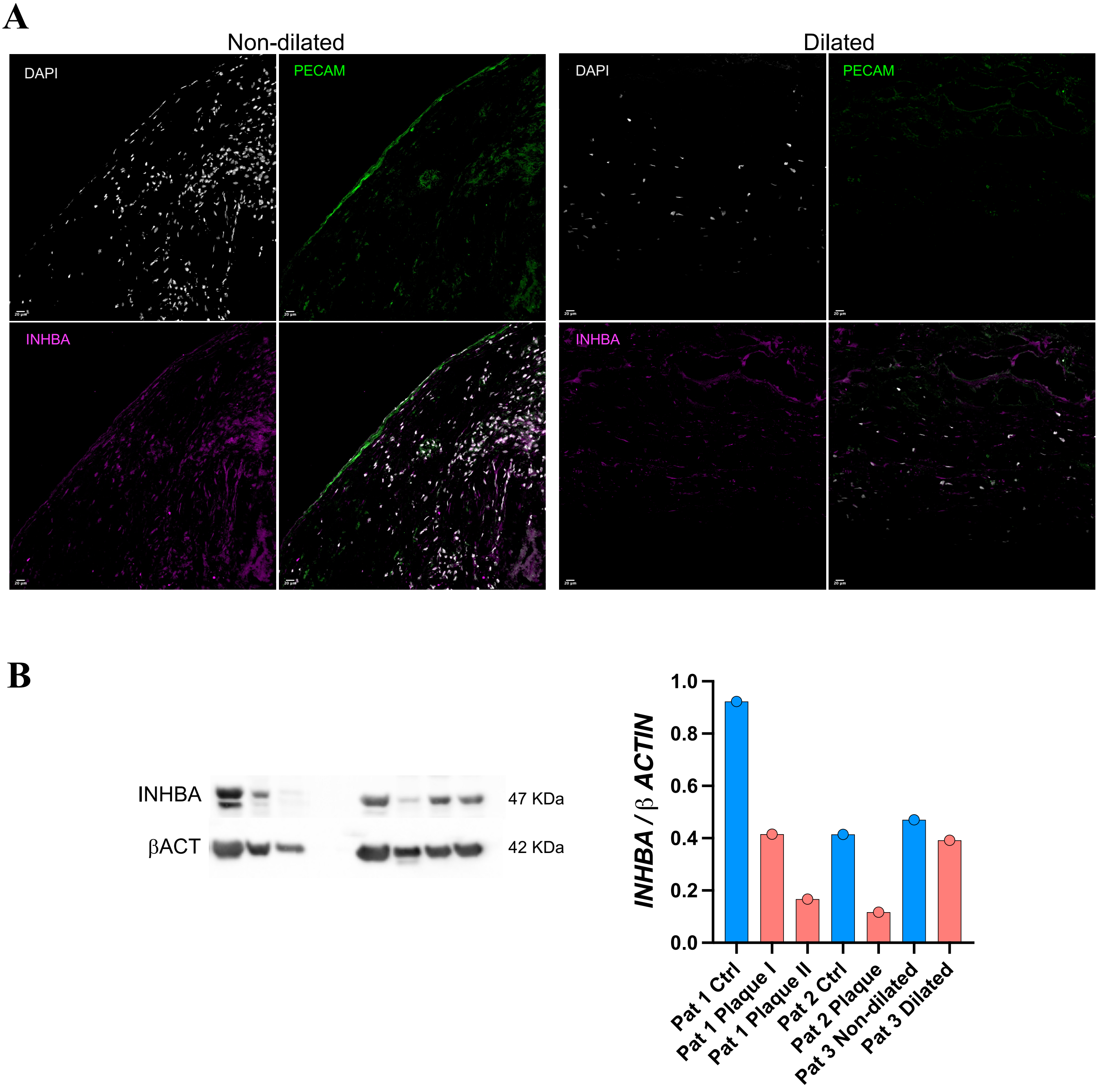


**Figure S4. Decrease amount of INHBA in aortas from AAA patinets and dieseased carotid (Plaque).** (**A**) Immunofluorescence on non-dilated and dilated aortic sections showing loss of INHBA (purple) as well as decrease degree co-localization with PECAM (green) signal in aneurysmal specimens. Imaging was carried out with confocal microscopy. (**B**) Protein lysates extracted from Ctrl/Plaque carotid tissues (n=2) and Non-dilated/Dilated aorta (n=1) were submitted to western blot analysis for the detection of INHBA. Protein levels are expressed as a ratio to beta actin (βACT). Quantification of the Western Blot was done with Fiji Image J software. Statistics: unpaired T-test (*) p-value $< 0.05$.

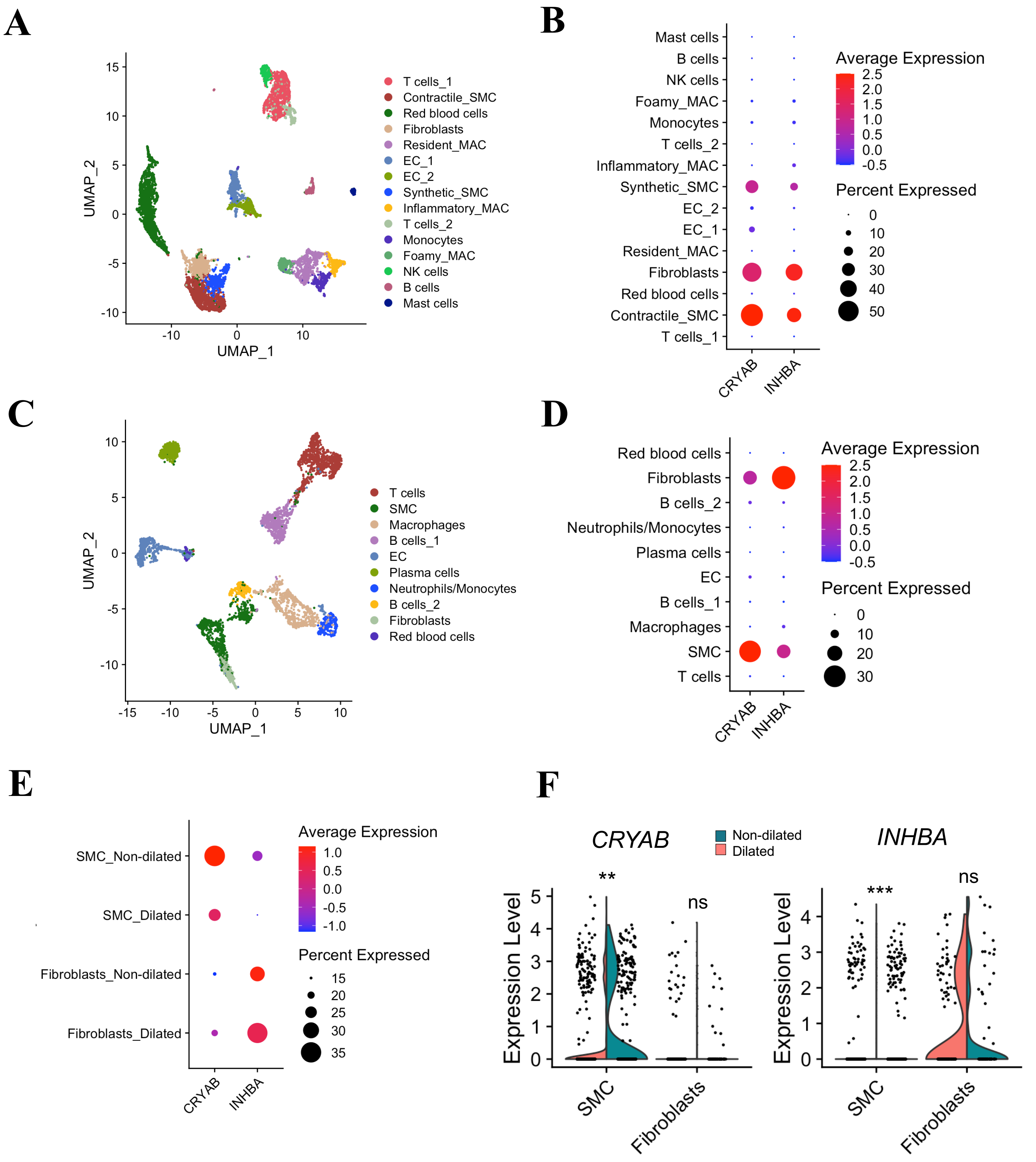


**Figure S5. ScRNAseq of human carotid palques and abdominal aortas.** (**A**) UMAP plot showing the major cells clusters identified from scRNAseq performed on human carotid plaques (n=5) and dot plot (**B**) showing the enrichment of *CRYAB* and *INHBA* in the "Fibro-SMC" cluster, highlighted in (**C**)UMAP plot showing the major cells clusters identified from scRNAseq performed on human AAA (n=3). (**D**) Dot plot showing the detection of *CRYAB* and *INHBA* in the "Fibro-SMC" cluster. (**E**) Dot and (**F**) violin plot showing the significantly enriched expression of *CRYAB* and *INHBA* in vascular SMC cluster from non-dilated aorta.

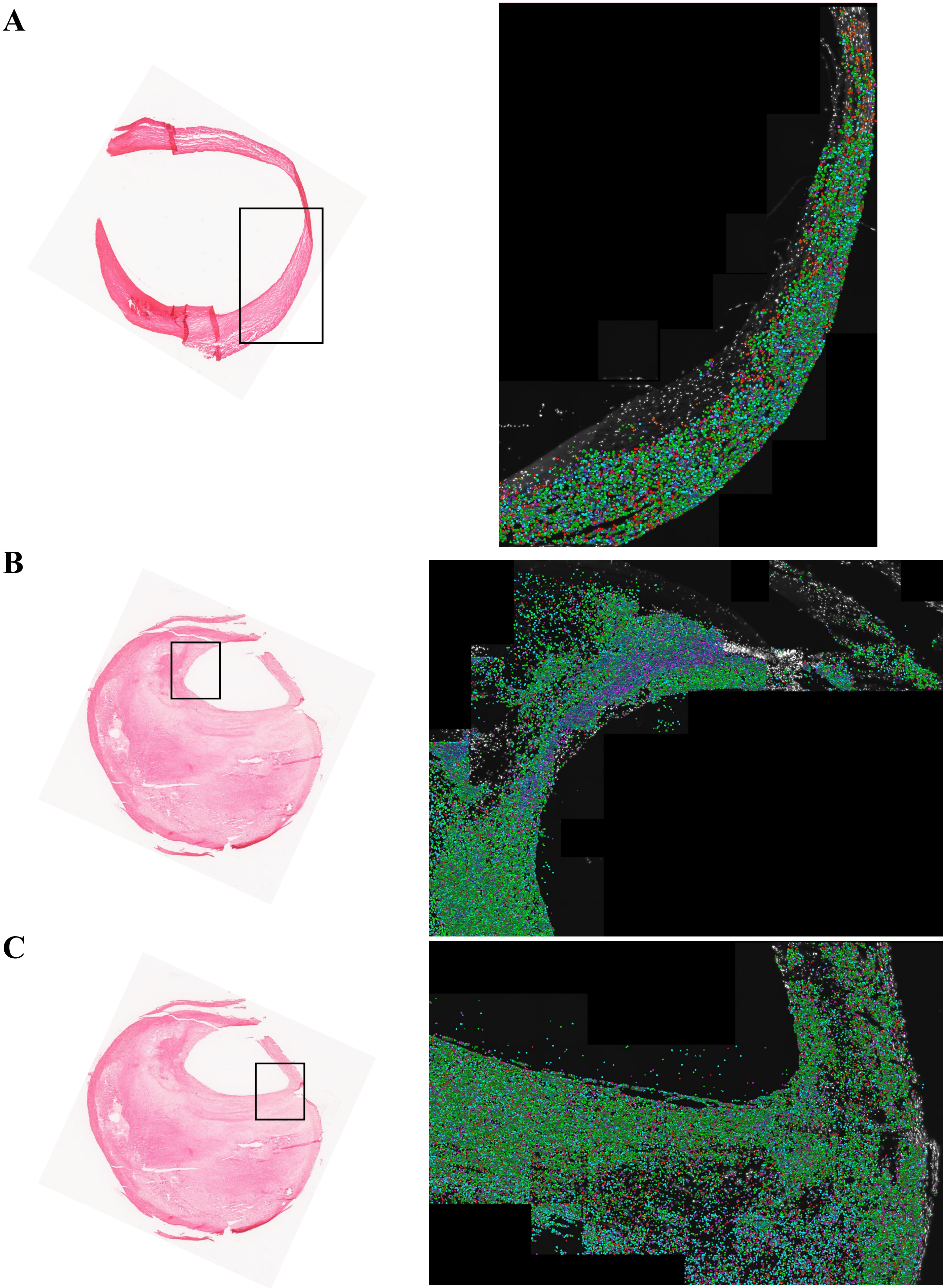


**Figure S6. Spatial transcriptomic of human carotid vessel and carotid plaque.** Ctrl (non-diseased carotid vessel) and carotid plaque were subjected to HybRISS methodology. All detected transcripts are shown: 16.583 transcripts in Ctrl (**A**), 78.245 and 75.920 in two different plaque regions respectively (**B**, **C**). The specific areas processed for spatial transcriptomic are indicated in the histochemistry images on the left side.

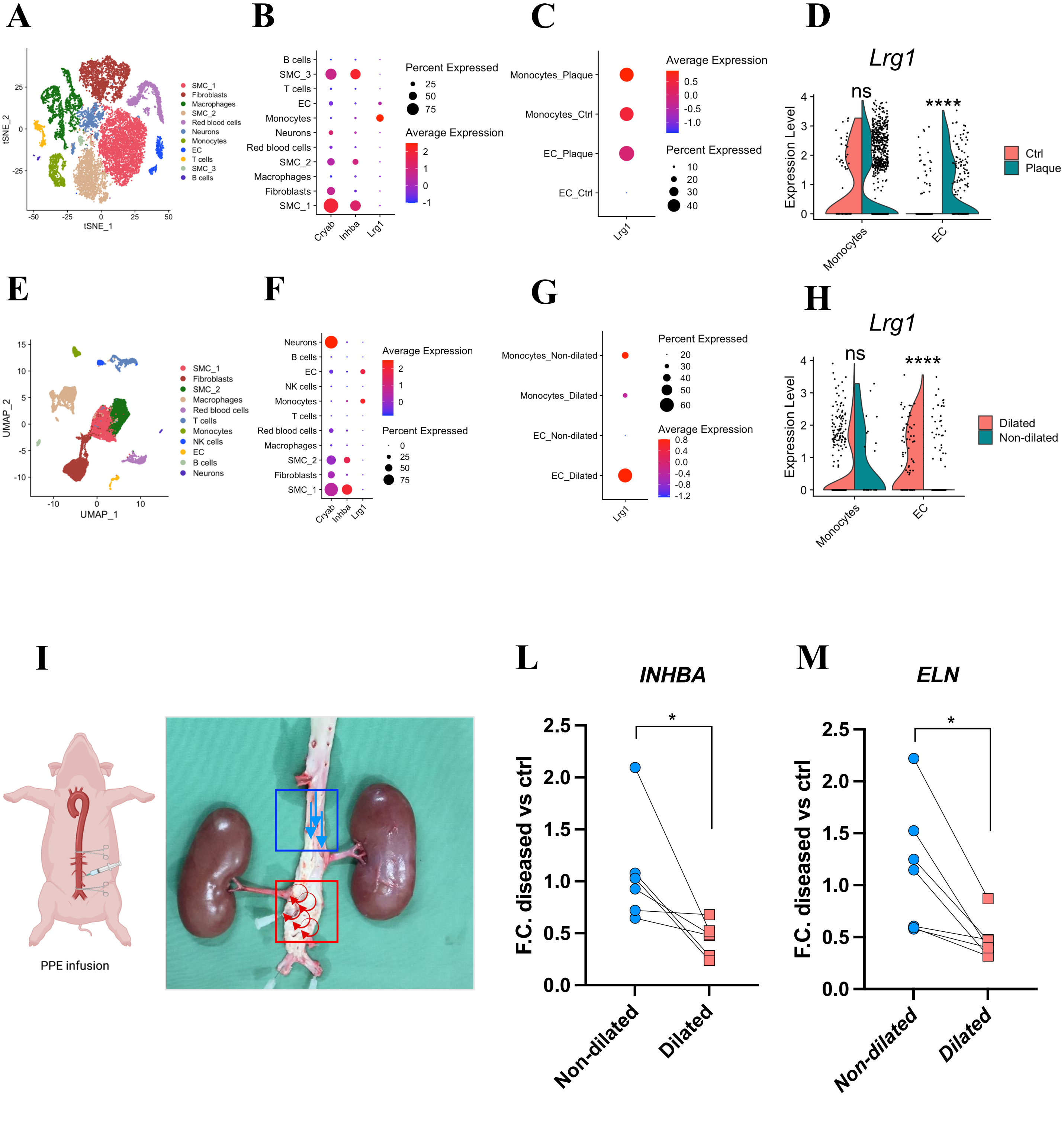


**Figure S7. *AoC*-generated targets in preclinical models of CAD and AAA.** (**A**) t-SNE plot showing the major cells clusters identified from scRNAseq of mice subjected to the carotid plaque rupture model (control tissues n=9, carotid plaque rupture tissues n=9). (**B**) The dot plot highlights the enrichment of *Lrg1* in Monocytes and EC clusters. (**C**) Dot and (**D**) violin plot show the significant enrichment of *Lrg1* in EC cluster originated from the diseased vessels. (**E**) t-SNE plot showing the major cells clusters identified via scRNAseq of mice subjected to the PPE-AAA model (control sham operated mice n=5, PPE infused mice n=6). (**F**) The dot plot on the right side highlights the enrichment of *Lrg1* in Monocytes and EC clusters. (**G**) Dot and (**H**) violin plot show the significant enrichment of *Lrg1* in EC cluster originated from AAA specimens. The FindMarkers function was used to compare the DEGs between the control and carotid plaques/AAA tissues, by using the default "Wilcoxon Rank Sum test". (****) p<0.0001, (***) p<0.0002, (**) p<0.0021, (*) p< 0.0332. (**I**) Picture of porcine PPE-AAA model. The biopsies were obtained from the aneurysm (red square) and the control non-dilated suprarenal segment (blue square). Blue arrows show the predicted laminar flow in non-dilated areas, rotating red arrows represents the predicted turbulent flow at the aneurysmatic areas. (**L, M**) QRT-PCR analysis of *INHBA* and *ELN* from non-dilated and dilated porcine aortas (n=7). Statistics: paired T-test (*) p-value < 0.05; (***) p-value < 0.001.

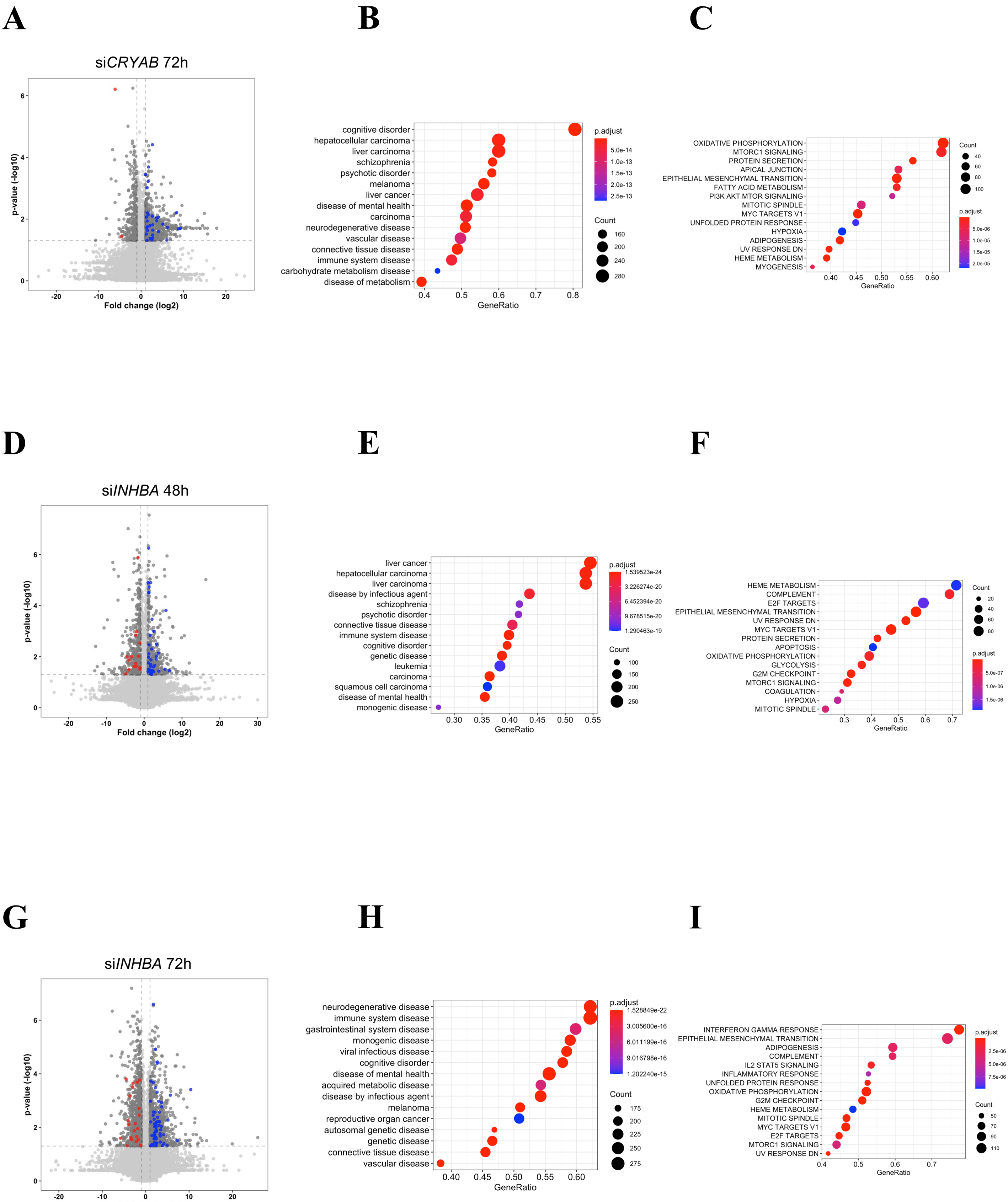


**Figure S8. Transcriptomic changes consequent to *CRYAB/INHBA* silencing in CaSMC.** **(A, D, G)** Volcano-plot with de-regulated genes 72h/48h after transfection with si*CRYAB*/si*INHBA* in CaSMC. Highlighted are genes contributing to GSEA score enrichment of Disease Ontology terms "Artery disease" and "Vascular disease". **(B, E, H)** Gene Set Enrichment Analysis (GSEA) using Disease Ontology and **(C, F, I)** MSigDB Hallmark gene sets.
